# Survival of hepatocytes from executioner caspase activation promotes liver regeneration by enhancing JAK/STAT3 activity

**DOI:** 10.1101/2024.08.28.610033

**Authors:** Zhiyuan Cao, Lining Qin, Kaixuan Liu, Chen Yao, Enhong Li, Xiaoyu Hao, Molin Wang, Baichun Jiang, Yongxin Zou, Huili Hu, Qiao Liu, Changshun Shao, Yaoqin Gong, Gongping Sun

**Affiliations:** Key Laboratory of Experimental Teratology, Ministry of Education, Department of Histology and Embryology, School of Basic Medical Sciences, Cheeloo College of Medicine, Shandong University, Jinan, Shandong, 250012, China; Key Laboratory of Experimental Teratology, Ministry of Education, Institute of Molecular Medicine and Genetics, School of Basic Medical Sciences, Cheeloo College of Medicine, Shandong University, Jinan, Shandong, 250012, China; Department of Systems Biomedicine and Research Center of Stem Cell and Regenerative Medicine, School of Basic Medical Sciences, Cheeloo College of Medicine, Shandong University, Jinan, Shandong, 250012, China; State Key Laboratory of Radiation Medicine and Protection, Institutes for Translational Medicine, Soochow University Suzhou Medical College, Suzhou, Jiangsu, 215123, China

## Abstract

Activation of executioner caspases, which is a key step in the apoptotic process, has been reported to promote tissue regeneration by sending pro-proliferation signals to the surrounding cells. However, whether executioner caspase activation (ECA) has cell-autonomous effect on tissue regeneration is not clear. Here, by generating transgenic mice carrying a lineage tracing system for cells that have experienced ECA, we demonstrate that transient ECA occurs in hepatocytes during liver regeneration after partial hepatectomy (PHx) or carbon tetrachloride (CCl_4_) treatment. Instead of committing apoptotic cell death, the majority of hepatocytes with ECA survive and proliferate to contribute to liver regeneration. Interestingly, inhibition of ECA in livers results in reduced hepatocyte proliferation and impaired regeneration, whereas increasing ECA to a level sufficient to kill hepatocytes also impedes regeneration, suggesting that ECA needs to be precisely controlled during liver regeneration. Mechanistic studies show that ECA promotes hepatocyte proliferation during regeneration through enhancing JAK/STAT3 activity. Our work reveals an essential role of survival of hepatocytes from ECA in liver regeneration.

## Introduction

Apoptosis is a conserved cell death program critical in organ development and homeostasis maintenance. Executioner caspases, which are caspase-3 and 7 in mammals, are key effectors in the apoptotic process whose activation disintegrates cells into apoptotic bodies^1, 2^. Apoptotic cells and executioner caspases have been reported as critical players in regeneration in diverse organisms including Hydra^3^, Drosophila^4^, zebrafish^5^, Xenopus^6^, salamander^7^ and mouse^8^. Upon injury, activated executioner caspases in apoptotic cells can cleave multiple protein substrates, leading to production and secretion of molecules like Wnt3^9^, PGE2^8^, EGFR ligands^10^, ATP^11^ to promote survival and proliferation of the neighboring cells. Activation of executioner caspases also leads to formation of apoptotic extracellular vesicles or apoptotic bodies^12–14^, which can be recognized by the neighboring cells or immune cells to initiate regenerative or reparative processes^15–17^. These reported regenerative effects exerted by executioner caspases are all cell non-autonomous, as it was long believed that upon injury, cells that activate executioner caspases would die.

In recent years, accumulating evidence has demonstrated that cells can survive executioner caspase activation (ECA) in response to stress^1, 18^. Wang et al. reported the existence of myofibers with active caspase-3 but no TUNEL staining in regenerating salamander limbs after amputation. Inhibition of caspase activity by overexpression of XIAP reduced dedifferentiation of myofibers after amputation^7^. Previously, using CasExpress, a lineage tracing system for cells that have experienced ECA, our group has shown that after X-ray radiation or transient overexpression of pro-apoptotic genes, a large group of cells in Drosophila wing imaginal discs can survive from ECA, proliferate and participate in formation of regenerated discs^19^. These studies suggest that ECA may have cell-autonomous effect on regeneration. Yet, whether survival from ECA occurs in regeneration of mammalian organs and how ECA in these survived cells regulates regeneration are not clear.

Liver is the most regenerative organ in adult mammals. After partial hepatectomy (PHx) or acute chemical injury, livers can restore the original weight and function^20^. To investigate the role of ECA in liver regeneration, we generated transgenic mice carrying mCasExpress reporter, the mammalian version of the CasExpress lineage tracing system. Using these mice, we demonstrate that ECA occurs at low frequency in the pericentral hepatocytes in the homeostatic liver and is dramatically upregulated during regeneration after PHx or carbon tetrachloride (CCl_4_) treatment. The ECA during liver regeneration is transient and preferentially takes place at the early stage. Of note, we show that hepatocytes that have experienced ECA need to survive to ensure rapid hepatocyte proliferation and liver regeneration, and that ECA promotes proliferation through increasing JAK-STAT3 activity.

## Results

### mCasExpress reporter reveals low frequency ECA in the homeostatic liver

To label cells that experience ECA in mice, we generated a transgenic mouse line carrying both *CAG-loxP-STOP-loxP-rtTA* (*LSL-rtTA*) and *TRE-Lyn11-NES-DEVD-FLP* and a transgenic mouse line carrying *CAG-FRT-STOP-FRT-ZsGreen* (*FSF-ZsGreen*). By crossing these two lines, we obtained the *LSL-rtTA; TRE-Lyn11-NES-DEVD-FLP; FSF-ZsGreen* mice, which were designated as *mCasExpress* mice. In *mCasExpress* mice, a fusion protein containing Lyn11 sequence, nuclear export signal (NES), an executioner caspase-specific cleavage site DEVD and a DNA recombinase FLP (LN-DEVD-FLP) is expressed in a Cre- and doxycycline (DOX)-dependent manner (Figure 1A). Without executioner caspase activity, FLP is tethered to the cell membrane. Once executioner caspases are activated, FLP is released from the membrane and translocate into the nucleus to remove the transcriptional termination signal (STOP) between the two FRT sites, leading to expression of the green fluorescent protein ZsGreen (Figure 1B). We crossed the *mCasExpress* mice with the *Sox2-Cre* mice, in which Cre is expressed in all the epiblast-derived tissues, to generate the *Sox2-Cre; mCasExpress* mice. Both the *mCasExpress* mice and the *Sox2-Cre; mCasExpress* mice exhibited normal appearance, body weight, liver weight, liver histology, serum aspartate transaminase (AST) and serum alanine transaminase (ALT) (Supplementary figure S1).

**Figure 1.**
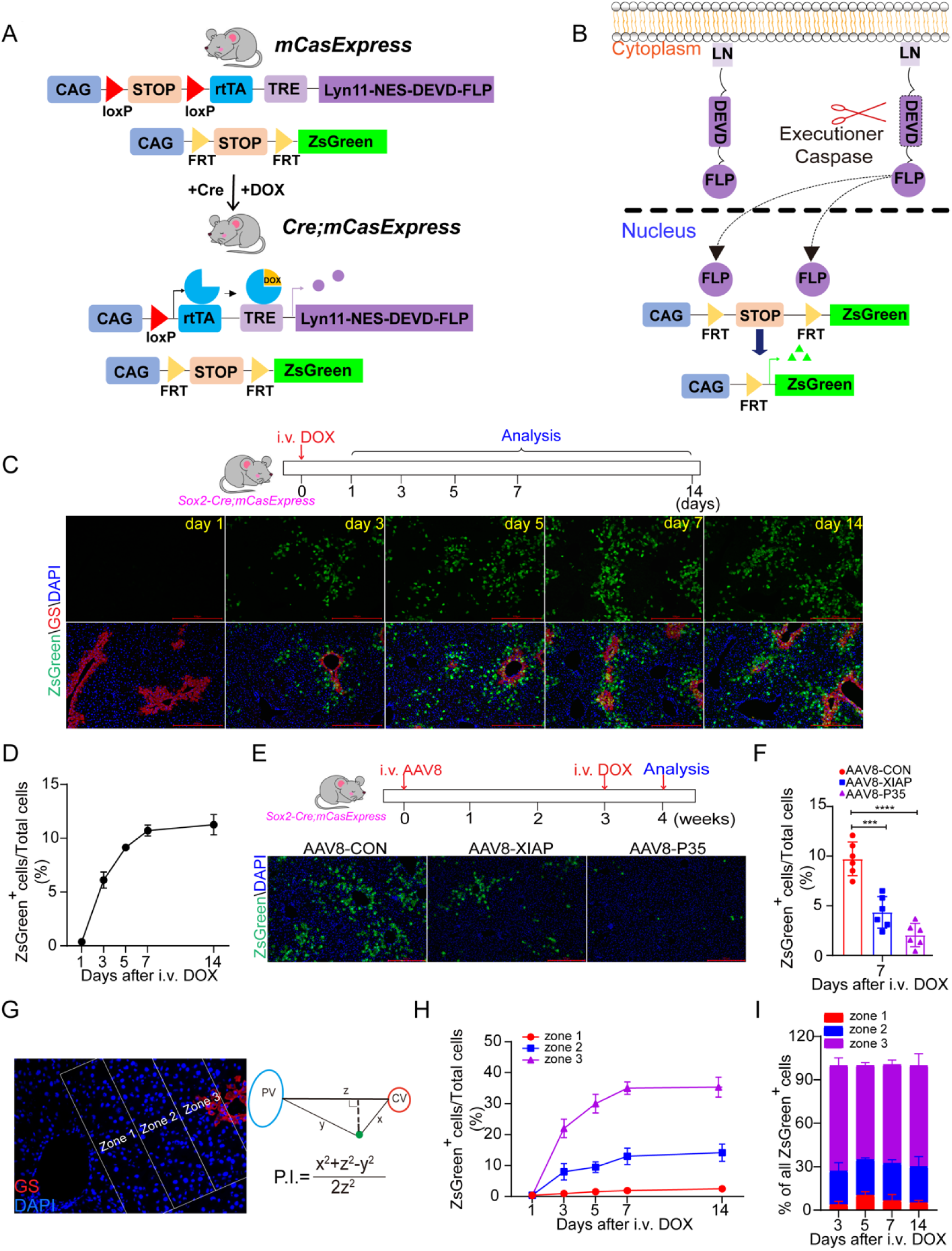
**ECA preferentially occurs in the pericentral region of the homeostatic liver.** A) The schematic of the mating strategy to generate *Cre; mCasExpress* mice. B) The schematic of mCasExpress. C) The representative images of livers at the indicated time points after injection of DOX. The staining for glutamine synthase (GS) marks the central veins. Scale bar: 100 μm. D) Quantification of the percentage of the ZsGreen^+^ cells in all cells within the field. For each time point, 3 mice were included and 3 fields per mouse were quantified. E) The representative images showing the effect of overexpressing XIAP or p35 on expression of ZsGreen in livers. Scale bar, 100 μm. F) Quantification of the percentage of the ZsGreen^+^ cells in total cells in the indicated groups. 6 mice per group and 3 fields per mouse. G) The zonation method. PV: portal vein. CV: central vein. P.I.: position index. Zone 1: P.I. ≥ 0.66. Zone 2: 0.33≤ P.I. <0.66. Zone 3: P.I.< 0.33. H) Quantification of the percentage of the ZsGreen^+^ cells in the total cells of each zone at the indicated time points. 3 mice per group and 3 fields per mouse. I) The bar graph shows the ratio of the number of the ZsGreen^+^ cells in zone 1, 2 or 3 to the number of all ZsGreen^+^ cells in the quantified lobule. 3 mice per group and 3 fields per mouse. i.v.: intravenous injection. Data are presented as the mean ± SD. ***: *P* <0.001. ****: *P* <0.0001.

Without DOX, LN-DEVD-FLP was rarely expressed and the *Sox2-Cre; mCasExpress* mice displayed no ZsGreen expression in the liver (Supplementary figure S2A & B). Transcription of *LN-DEVD-FLP* was highly induced after injection of 5 mg/kg DOX through tail veins and diminished within 7 days after injection (Supplementary figure S2A). We assessed the ZsGreen expression in the livers from the *Sox2-Cre; mCasExpress* mice after injection of DOX. Few ZsGreen^+^ cells were detected on day 1 after injection (Figure 1C & D), possibly because one day was insufficient for cells to accumulate a detectable level of ZsGreen protein. The percentage of the ZsGreen^+^ cells increased to 6% on day 3 after injection and 10.7% on day 7 (Figure 1C & D). From day 7 to day 14, the percentage of the ZsGreen^+^ cells stayed constant (Figure 1C & D), in consistence with the diminished LN-DEVD-FLP expression (Supplementary figure S2A). The *mCasExpress* mice or the *Sox2-Cre; FSF-ZsGreen* mice contained no ZsGreen^+^ cells in livers after DOX injection (Supplementary figure S2C), supporting the requirement of FLP activity in ZsGreen expression. The ZsGreen^+^ cells were also observed in the homeostatic livers after DOX injection when *mCasExpress* was driven by the liver-specific *Alb-Cre* (Supplementary figure S3). To determine whether the expression of ZsGreen relies on caspase activity, the *Sox2-Cre; mCasExpress* mice were administered with adeno-associated virus serotype 8 (AAV8) expressing baculovirus p35 or mouse XIAP, both of which are inhibitors of caspases^21^. Overexpression of either p35 or XIAP dramatically reduced the percentage of the ZsGreen^+^ cells in livers (Figure 1E & F), indicating that the ZsGreen^+^ cells are cells that have experienced caspase activation.

### ECA occurs preferentially in the pericentral hepatocytes of the homeostatic liver

To determine the cell types that activate executioner caspases in liver, we stained the *Sox2-Cre; mCasExpress* liver samples collected on day 7 after DOX injection with markers of the major cell types in the liver. We found that all the ZsGreen^+^ cells in liver expressed the hepatocyte marker HNF4α, and detected no co-localization of ZsGreen with the cholangiocyte marker CK19, the macrophage marker CD68 or the endothelial cell marker CD31 (Supplementary figure S4), suggesting that ECA only occurs in hepatocytes. Hepatocytes are highly heterogenous^22^. We analyzed the spatial distribution of the ZsGreen^+^ hepatocytes using the method reported by Wei et al.^23^, in which the position index (P.I.) of a hepatocyte was calculated based on its distance to the closest central vein, which was surrounded by GS (glutamine synthase)^+^ hepatocytes, and the distance to the closest portal vein (Figure 1G). In the homeostatic liver after DOX injection, the percentage of the ZsGreen^+^ hepatocytes increased much faster in the pericentral zone 3 (P.I. < 0.33) than in the other two zones (Figure 1H). At all the time points we analyzed, about 70 % of the ZsGreen^+^ cells were in zone 3 while less than 10 % of the ZsGreen^+^ cells were in zone 1 (P.I. > 0.66) (Figure 1I). To figure out whether the pericentral enrichment of the ZsGreen^+^ hepatocytes are due to uneven distribution of ECA or DOX, we injected DOX to the *CAG-rtTA; tetO-Cre; LSL-tdTomato* mice, in which Cre expression is induced by DOX and leads to permanent tdTomato expression. On day 7 after injection, the tdTomato^+^ cells were uniformly distributed in the liver (Supplementary figure S5). Therefore, the pericentral enrichment of the ZsGreen^+^ hepatocytes indicates a higher frequency of ECA in the pericentral region.

### Transient ECA is induced in the early stage of liver regeneration

To determine whether executioner caspases are activated during liver regeneration, we performed 70% PHx on the *Sox2-Cre; mCasExpress* mice at 24 hours after DOX injection and harvested the livers on day 1, 3, 5 and 7 after PHx. As reported in literature, the liver/body weight ratio was mostly restored to the control level on day 7 (Figure 2A). The levels of serum AST and ALT were restored on day 3 (Figure 2B & C). On day 1 after surgery, livers from the PHx group contained more ZsGreen^+^ cells than those from the sham group. The percentage of ZsGreen^+^ cells in the regenerating livers strongly increased from day 1 to day 3 and exhibited dramatic difference compared to the sham group. On day 7, the livers from the PHx group contained about 30% ZsGreen^+^ cells, while only 10% cells in the livers from the sham group expressed ZsGreen (Figure 2D & E). Overexpression of p35 or XIAP significantly suppressed the ZsGreen expression after PHx (Figure 2F & G), indicating that the ZsGreen signal was a consequence of caspase activation. The elevated ECA was also observed in the regenerated livers from the *Alb-Cre; mCasExpress* mice and the *CAG-Cre; mCasExpress* mice after PHx (Supplementary figure S6).

**Figure 2.**
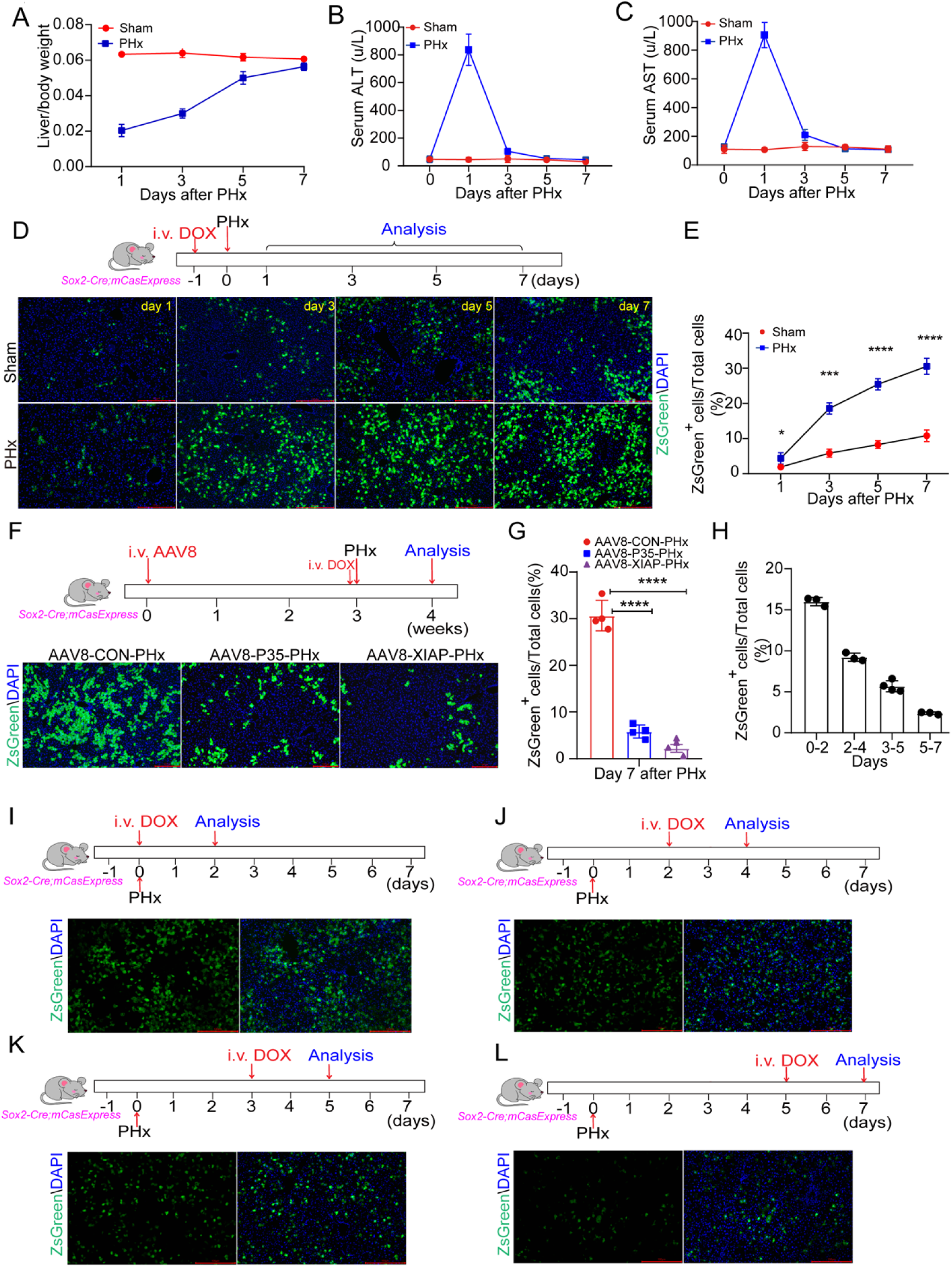
**Transient ECA is induced during regeneration after PHx.** A-C) Change of the ratio between liver weight and body weight (A), serum ALT (B) and serum AST (C) within 7 days after PHx. 3 mice in each group. D-E) The representative images and quantification of the ZsGreen^+^ cells in the livers at different time points after sham operation or PHx. Scale bar: 100 μm. 5 mice per group and 3 fields per mouse. F-G) The representative images and quantification of the ZsGreen^+^ cells showing the effect of XIAP or p35 overexpression on ZsGreen expression. Scale bar: 100 μm. 4 mice per group and 3 fields per mouse. H-L) Analysis of the ZsGreen^+^ cells shown up within a two-day window during regeneration after PHx. DOX was injected on day 0 (I), day 2 (J), day 3 (K), day 5 (L) and livers were collected two days after DOX injection. i.v.: intravenous injection. Data are presented as the mean ± SD. *: *P* <0.05. **: *P* <0.01. ***: *P* <0.001. ****: *P* <0.0001.

In Figure 2D-E, the 30% ZsGreen^+^ hepatocytes detected on day 7 were the sum of hepatocytes that experienced ECA and their descendants over 7-day regeneration. To monitor the dynamics of ECA in the process of regeneration, we injected DOX at different time points after PHx and collected liver samples at 48 hours post-injection. We found that about 16% hepatocytes became ZsGreen^+^ between day 0 and day 2 after PHx, and the percentage reduced to 9% between day 2 and day 4 and to 5.5% between day 3 and day 5. In the terminating phase of regeneration (from day 5 to day 7), only 2.5% of hepatocytes were ZsGreen^+^ (Figure 2H-L). TUNEL assays revealed that the level of apoptotic cell death during regeneration was very low, especially in the first three days (Supplementary figure S7), suggesting that most of the hepatocytes with ECA survived. These data indicate that ECA during regeneration is transient and occurs more frequently in the early stage.

To determine whether the elevated fraction of hepatocytes that have survived from ECA is specific to the regenerated liver after PHx, we peritoneally injected 10% CCl_4_ to the *Sox2-Cre; mCasExpress* mice and the *CAG-Cre; mCasExpress* mice to induce acute liver injury. Injection of 10% CCl_4_ induced strong increase in the ZsGreen^+^ cells in livers after 7-day regeneration (Supplementary figure S8), suggesting survival from ECA may be a common process involved in liver regeneration.

We then analyzed the distribution of the ZsGreen^+^ cells in the regenerated livers. On day 7 after PHx or CCl_4_ injection, the highest percentage of the ZsGreen^+^ hepatocytes was observed in zone 2 (Figure 3A & B, D & E). ZsGreen^+^ cells were also observed in zone 1, but were largely excluded in the region next to the portal vein (Figure 3C & F), suggesting the resistance of these hepatocytes to ECA. The difference in the percentage of ZsGreen^+^ cells between the regenerated liver and the sham control was the lowest in zone 3, possibly due to the accumulation of ZsGreen^+^ cells in the pericentral region over 7 days in the sham group. Thus, we analyzed the distribution of the ZsGreen^+^ cells on day 2 after surgery when the percentage of ZsGreen^+^ cells in the sham group was low. The ZsGreen^+^ cells were strongly increased in all three zones after PHx (Figure 3G & H).

**Figure 3.**
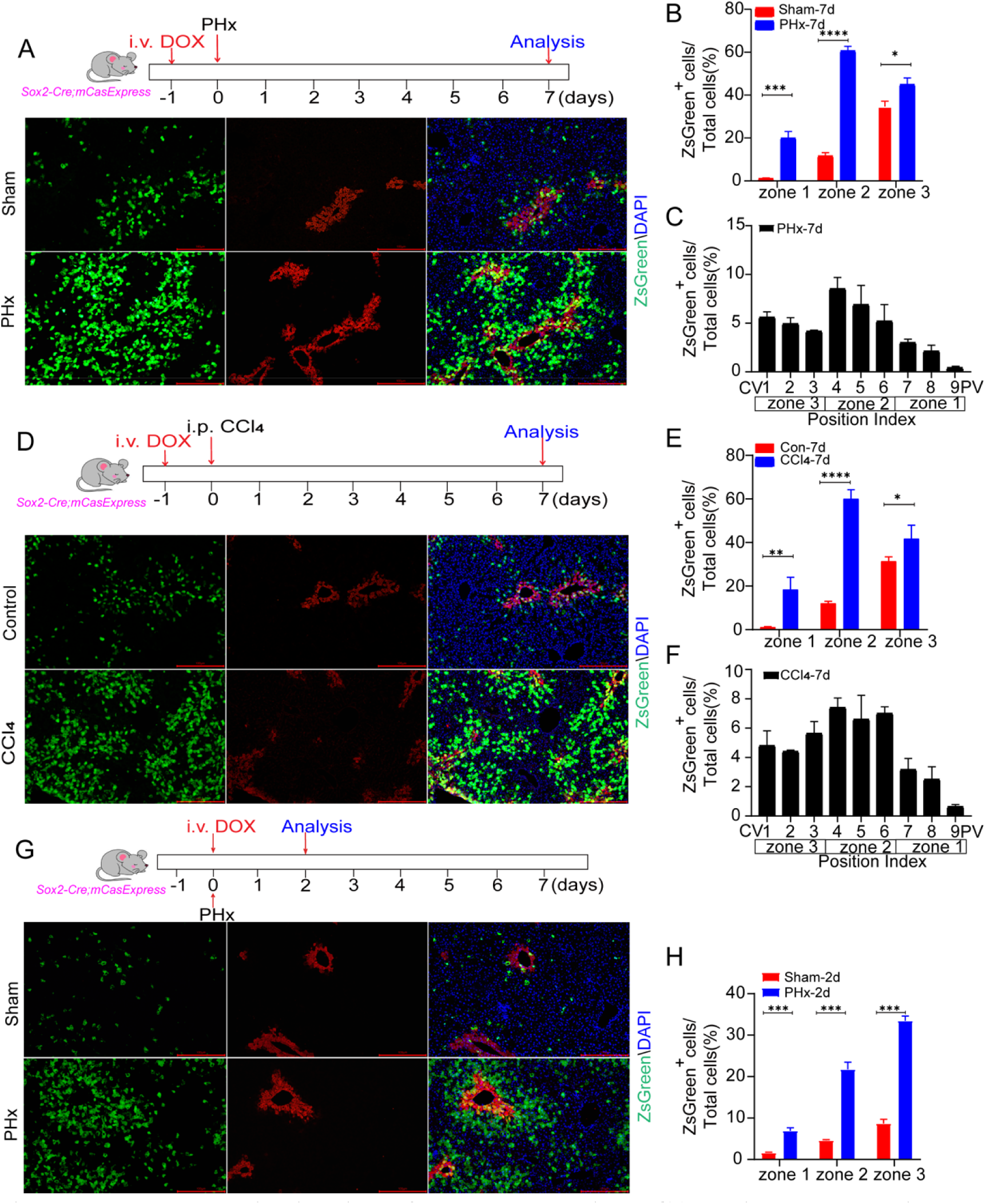
**The zonal distribution of hepatocytes with ECA during regeneration.** A) The representative images of livers on day 7 after PHx or sham operation with staining of pericentral region marker GS. Scale bar: 100 μm. DOX was injected one day before PHx. B) Quantification of the percentage of the ZsGreen^+^ cells in the total cells of each zone on the samples shown in (A). C) The percentage of the ZsGreen^+^ cells in each position of the liver lobule shown in (A). 3 mice per group and 3 fields per mouse. D) The representative images of livers on day 7 after injection of CCl_4_ or corn oil (control) with staining of pericentral region marker GS. Scale bar: 100 μm. DOX was injected one day before CCl_4_ injection. i.p. intraperitoneal injection. E) Quantification of the percentage of the ZsGreen^+^ cells in the total cells of each zone on the samples shown in (D). F) The percentage of the ZsGreen^+^ cells in each position of the liver lobule shown in (D). 4 mice in the CCl_4_ group and 3 mice in the control group. 3 fields per mouse. G) The representative images of livers on day 2 after PHx with staining of pericentral region marker GS. Scale bar: 100 μm. DOX was injected on the day of PHx. H) Quantification of the percentage of the ZsGreen^+^ cells in the total cells of each zone on the samples shown in (G). Data are presented as the mean ± SD. *: *P* <0.05. **: *P* <0.01. ***: *P* <0.001. ****: *P* <0.0001.

### ECA is required for hepatocyte proliferation and liver regeneration

To investigate the role of ECA in liver regeneration, we first inhibited caspase activity in livers by overexpressing XIAP or p35. Without injury, neither the liver/body weight ratio nor the serum ALT and AST levels were affected by overexpression of XIAP or p35 (Supplementary figure S9A & B). However, the restoration of the liver/body weight ratio after PHx was significantly impaired in livers overexpressing XIAP or p35 (Figure 4A). Regeneration after PHx relies on hepatocyte proliferation. We therefore evaluated the effect of caspase inhibition on hepatocyte proliferation. The XIAP or p35-overexpressing livers collected on day 2 after PHx contained fewer Ki67^+^ hepatocytes and lower levels of Cyclin D1 and Cyclin E1 than the control livers (Figure 4B-E), suggesting that inhibition of caspases suppresses hepatocyte proliferation. To further confirm the role of executioner caspases in hepatocyte proliferation and liver regeneration, we generated mice with liver-specific knockout of both *Casp3* and *Casp7* (designated as DKO mice), which encode caspase-3 and 7, respectively. DKO mice exhibited liver/body weight ratio and serum AST and ALT similar to the *Casp3^flox/flox^ Casp7^flox/flox^*littermates (designated as CON mice) (Supplementary figure S9C-E). Knockout of both *Casp3* and *Casp7* inhibited restoration of liver/body weight ratio and hepatocyte proliferation after PHx (Figure 4F-J). Furthermore, we found that the ZsGreen^+^ hepatocytes had a higher Ki67^+^ fraction than the ZsGreen^-^ hepatocytes from the same liver (Figure 4K-L). These data together indicate that executioner caspases promote hepatocyte proliferation during liver regeneration.

**Figure 4.**
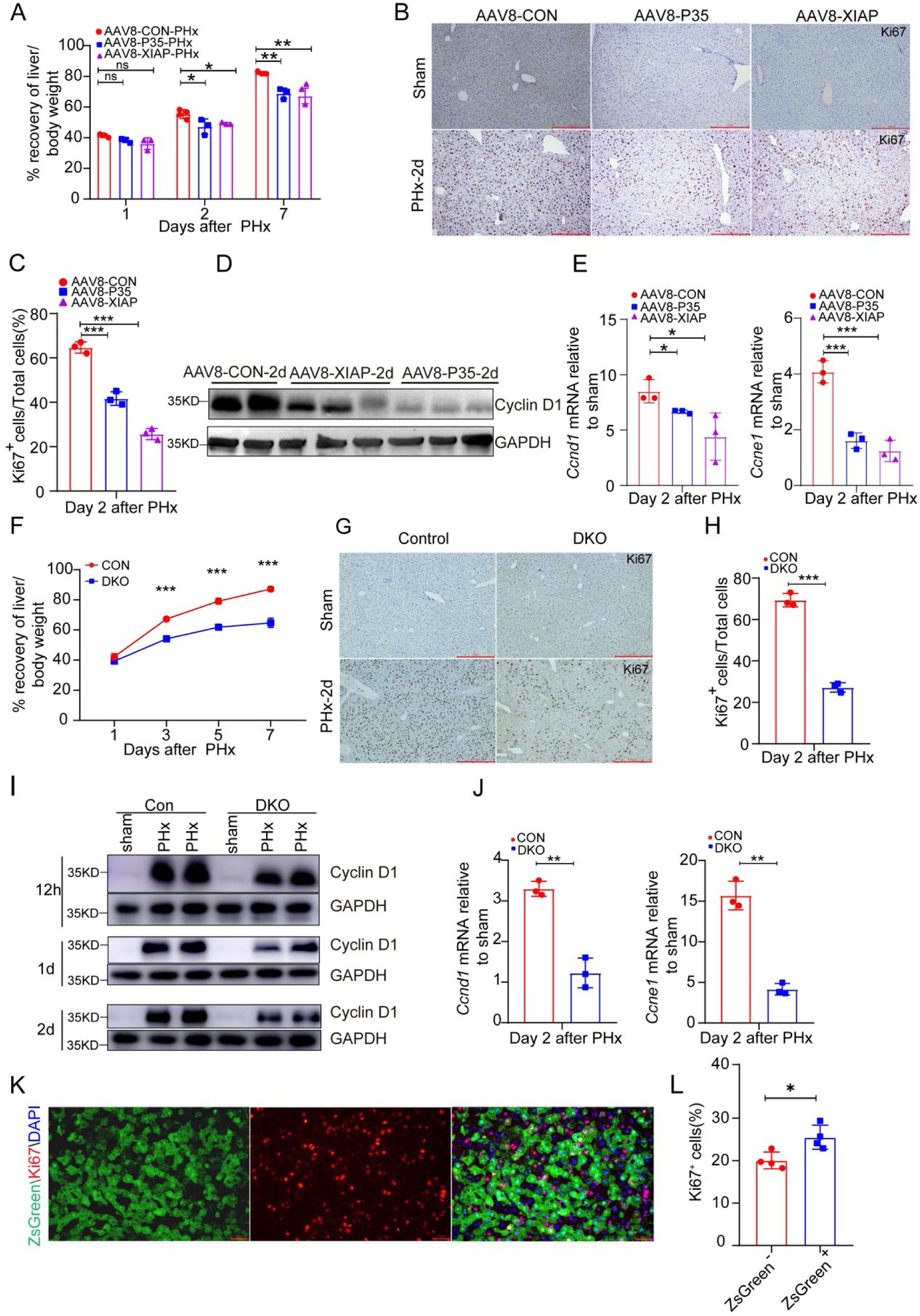
**Inhibition of executioner caspase activity or expression impairs liver regeneration after PHx.** A) The percentage of recovery of the liver/body weight ratio at the indicated time points after PHx. The average liver/body weight ratio of the sham group in each genotype was considered as 100%. N=3 for all groups except the AAV8-CON-PHx on day 2, for which N=4. B) The representative images of Ki67 staining in the indicated groups on day 2 after sham operation or PHx. Scale bar: 100 μm. C) Quantification of the percentage of Ki67^+^ cells in the indicated groups on day 2 after PHx. 3 mice per group and 3 fields per mice. D) Western blots showing the protein level of Cyclin D1 in the indicated groups on day 2 after PHx. E) The mRNA levels of *Ccnd1* and *Ccne1* in the indicated groups on day 2 after PHx. The level in the sham operated animals was set as 1. 3 mice per group. F) The percentage of recovery of the liver/body weight ratio after PHx in *Casp3^fl/fl^ Casp7^fl/fl^* (CON) mice and *Alb-Cre^+/-^ Casp3^fl/fl^ Casp7^fl/fl^*(DKO) mice. The average liver/body weight ratio of the sham group in each genotype was considered as 100%. 3 mice per group. G-H) The representative images and quantification of Ki67 staining on day 2 after surgery in the indicated groups. Scale bar: 100 μm. 3 mice per group and 3 fields per mouse. I) Wester blots showing Cyclin D1 level at 12 hr, 1 day and 2 day after surgery in the indicated groups. J) The mRNA levels of *Ccnd1* and *Ccne1* in the indicated livers. 3 mice per group. K) The representative images showing the co-localization of ZsGreen and Ki67 in the liver on day 3 after PHx. Scale bar: 50 μm. L) Quantification of the Ki67^+^ cells in the ZsGreen^+^ and the ZsGreen^-^ cells on day 3 after PHx. 4 mice per group and 3 fields per mouse. Data are presented as the mean ± SD. *: *P* <0.05. **: *P* <0.01. ***: *P* <0.001. ns: no significance.

### Survival of hepatocytes with ECA is necessary for liver regeneration

An important function of ECA is to execution apoptotic cell death. Apoptosis can drive compensatory proliferation of the surrounding cells, which has been reported to contribute to tissue regeneration in diverse organisms like Drosophila, Hydra, Xenopus^3, 4, 6^. However, the facts that many ZsGreen^+^ hepatocytes can proliferate and that apoptotic cells were rarely detected in the early stage of regeneration suggest that ECA drives hepatocyte proliferation independent of apoptosis execution. To verify this hypothesis, we developed a genetic ablation system consisting of the executioner caspase-activatable FLP (LN-NES-DEVD-FLP) and the CAG-FRT-STOP-FRT-tBid cassette, which can express tBid after removal of the transcription termination signal (STOP) by FLP. In cells carrying this system, activation of executioner caspases can induce overexpression of tBid, the cleaved form of the BH3-only protein Bid that can trigger activation of apoptotic caspase cascade by inducing MOMP^24, 25^, leading to rapid amplification of ECA and commitment of cell death (Figure 5A). If the hepatocytes with ECA eventually die and promote liver regeneration through apoptosis-induced proliferation, allowing these cells to die will have no or even positive effect on liver regeneration. If survival of hepatocytes with ECA is necessary for regeneration, their death will impair liver regeneration.

**Figure 5.**
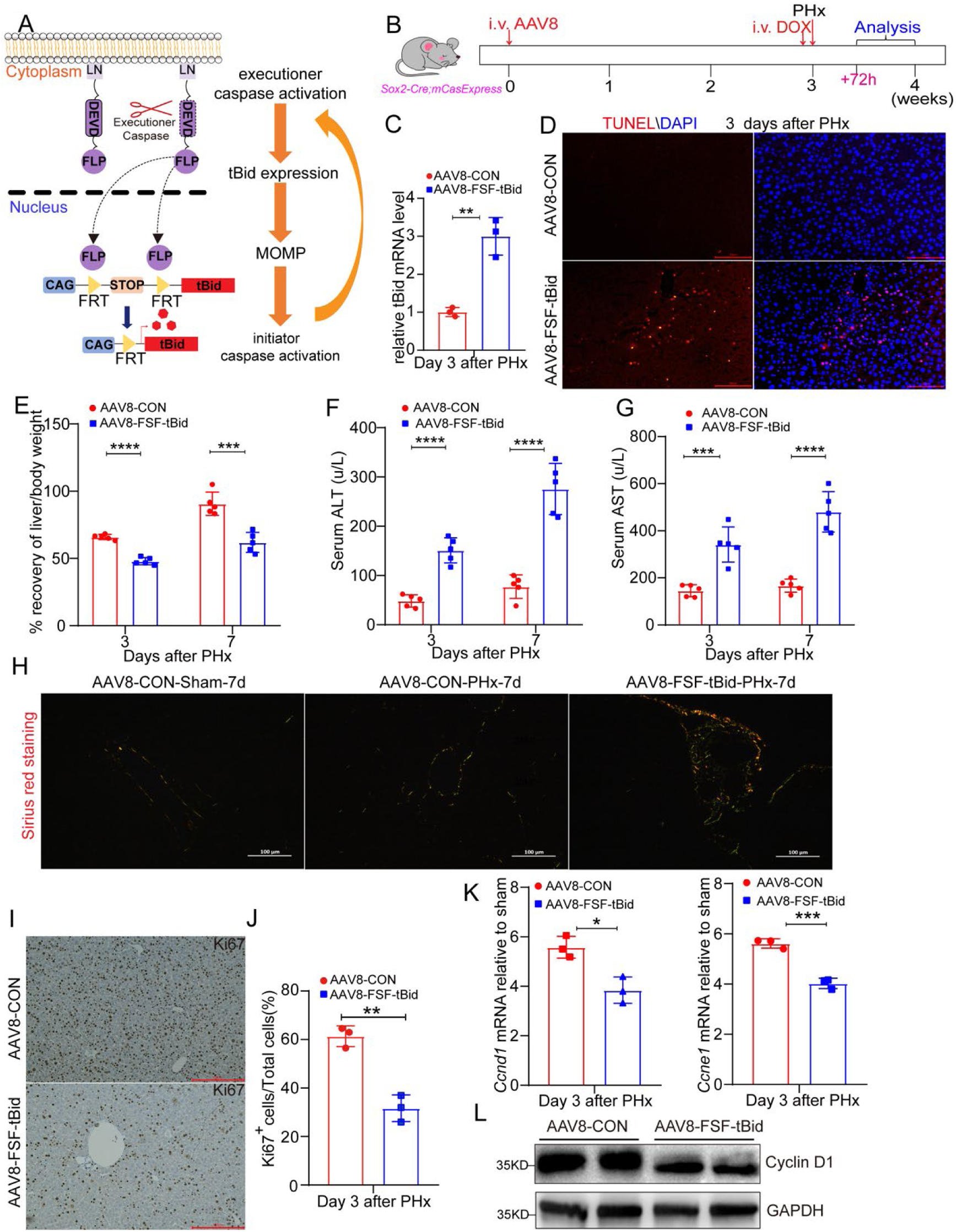
**Death of hepatocytes with ECA suppresses liver regeneration.** A) The schematic of ablating cells with ECA. B) The workflow of experiments in this figure. i.v.: intravenous injection. C) The mRNA expression of *tBid* in livers on day 3 after PHx. 3 mice per group. D) The representative images of TUNEL assays in livers on day 3 after PHx. Scale bar: 50 μm. E) The percentage of recovery of the liver/body weight ratio at the indicated time points after PHx. The average liver/body weight ratio of the sham group was considered as 100%. 5 mice per group. F-G) The serum ALT (F) and AST (G) at the indicated time points after PHx. 5 mice per group. H) Pricosirius red staining of the indicated livers on day 7 after PHx. The images were captured under polarized light. Scale bar: 100 μm. I-J) The representative images and quantification of Ki67 staining in the indicated livers on day 3 after PHx. Scale bar: 100 μm. 3 mice per group and 3 fields per mouse. K) The mRNA levels of *Ccnd1* and *Ccne1* in the indicated livers on day 3 after PHx. 3 mice per group. L) Western blots showing Cyclin D1 level the indicated livers on day 3 after PHx. Data are presented as the mean ± SD. *: *P* <0.05. **: *P* <0.01. ***: *P* <0.001. ****: *P* <0.0001.

We infected the *Sox2-Cre; mCasExpress* mice with AAV8 carrying *CAG-FRT-STOP-FRT-tBid* (AAV8-FSF-tBid). Without DOX, the *Sox2-Cre; mCasExpress* mice injected with AAV8-FSF-tBid displayed body weight and liver weight similar to those administered with control AAV8 (AAV8-CON) (Supplementary figure S10A). 7 days after DOX injection, the *Sox2-Cre; mCasExpress* mice injected with AAV8-FSF-tBid exhibited mildly increased serum ALT and AST compared to those with AAV8-CON (Supplementary figure S10B), possibly due to ablation of the hepatocytes with ECA under homeostasis. We then performed PHx on the *Sox2-Cre; mCasExpress* mice administered with AAV8-FSF-tBid or AAV8-CON at 24 hour after DOX injection (Figure 5B). Three days after PHx, compared to the livers injected with AAV8-CON, those with AAV8-FSF-tBid expressed a higher level of tBid (Figure 5C) and contained more TUNEL^+^ apoptotic cells (Figure 5D), indicating that induction of tBid overexpression in cells with ECA did drive more cell death. On day 3 and day 7 after PHx, mice with AAV8-FSF-tBid exhibited significantly lower restoration of the liver/body weight ratio and markedly higher AST and ALT (Figure 5E-G). Sirius red staining revealed elevated collagen deposition in the liver with AAV8-FSF-tBid after 7-day regeneration (Figure 5H). We then assessed the effect of cell death on hepatocyte proliferation in early regeneration. Compared to livers with AAV8-CON, the livers with AAV8-FSF-tBid exhibited a reduced frequency of Ki67^+^ hepatocytes (Figure 5I & J) and downregulated expression of Cyclin D1 and Cyclin E1 (Figure 5K & L), indicative of suppressed proliferation. These data together suggest that survival of cells with ECA is required to ensure rapid hepatocyte proliferation and proper liver regeneration.

### ECA promotes hepatocyte proliferation through enhancing activity of JAK/STAT3 signaling

The next question is how ECA promotes hepatocyte proliferation. We observed reduced Cyclin D, which is critical in driving hepatocyte into S phase, when executioner caspases were inhibited or depleted (Figure 4D & I). During liver regeneration, expression of Cyclin D is mainly induced by STAT3 activation. Loss of STAT3 inhibits Cyclin D expression and hepatocyte proliferation while overexpression of the constitutively active STAT3 in hepatocytes increases proliferation^26–28^. Therefore, we assessed STAT3 activity in livers overexpressing XIAP or p35. STAT3 phosphorylation and expression of the target gene *Socs1* were dramatically inhibited in the livers overexpressing XIAP or p35 on day 1 and 2 after PHx (Figure 6A & B), suggesting reduced STAT3 activation. Consistently, suppressed STAT3 activation in the early stage of regeneration after PHx was also detected in livers with AAV8-FSF-tBid (Figure 6C & D) and in DKO livers (Figure 6E & F), suggesting that executioner caspases promote STAT3 activation during liver regeneration. Recently, Zhu et al. reported that caspase-3 induces STAT3 phosphorylation through DNA damage-induced activation of the non-receptor protein tyrosine kinase Src during oncogenic transformation^29^. However, little DNA damage was detected in regenerating livers after PHx (Supplementary figure S11), suggesting that executioner caspases promote STAT3 activation through a different mechanism. In liver regeneration, STAT3 is known to be activated by IL-6^30^. IL-6 binds to its receptor to activate JAK family protein, leading to phosphorylation of STAT3^31^. Hence, we tested IL-6 level and phosphorylation of JAK2 in CON and DKO mice after PHx. Knockout of both *Casp3* and *Casp7* reduced phosphorylation of JAK2 (Figure 6E). However, the serum levels of IL-6 in CON and DKO mice after PHx were similar (Figure 6G). Moreover, primary hepatocytes isolated from DKO livers displayed reduced phosphorylation of STAT3 and JAK2 in response to IL-6 treatment (Figure 6H). These data suggest that executioner caspases promote STAT3 activation downstream of IL-6 and upstream of JAK.

**Figure 6.**
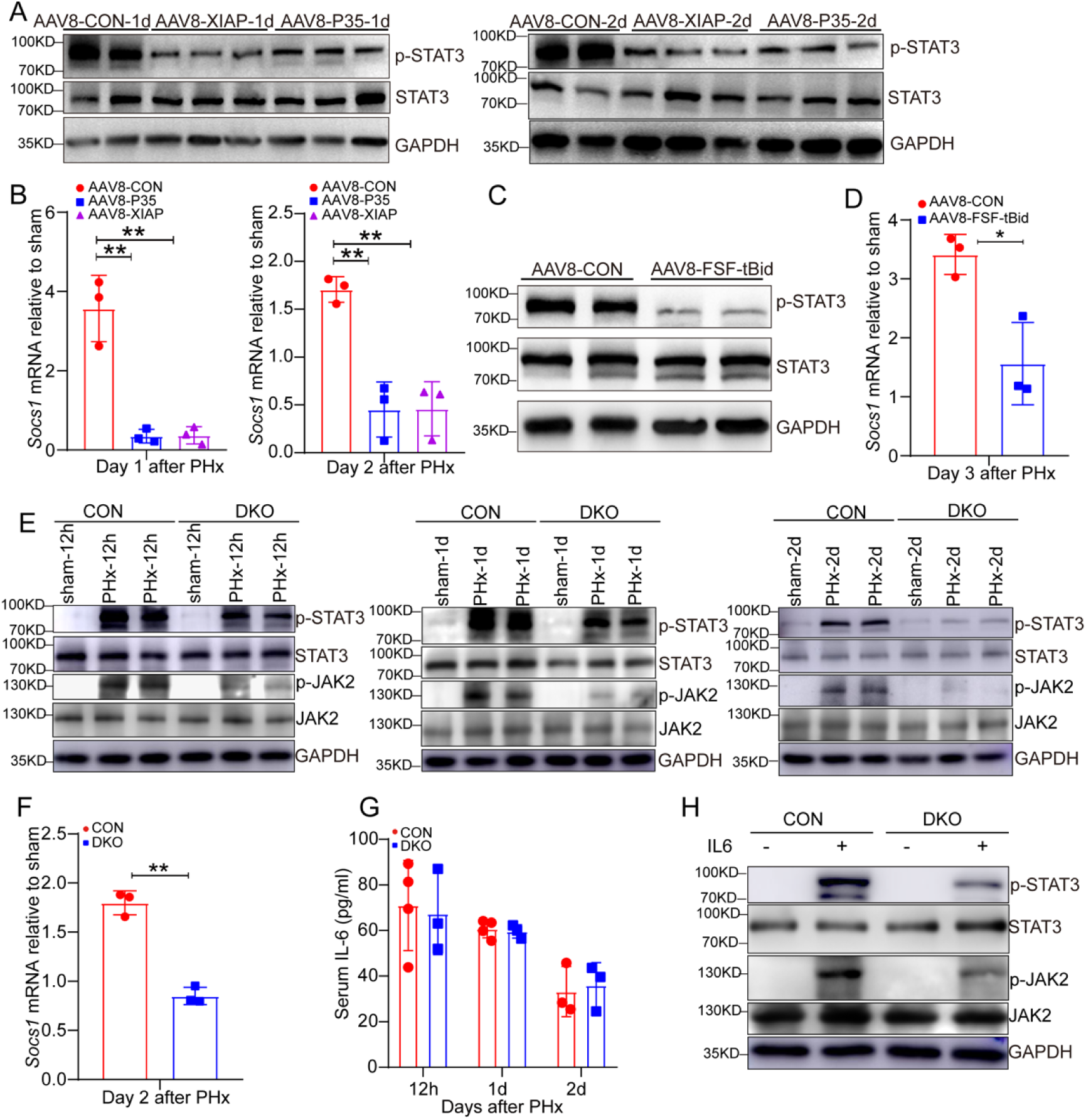
**ECA increases JAK/STAT3 signaling activity.** A) Western blots showing the protein levels of p-STAT3 and STAT3 in livers administered with the indicated AAV8 on day 1 and 2 after PHx. B) The mRNA level of *Socs1* in the indicated groups on day 1 and 2 after PHx. The level in the sham operated animals was set as 1. 3 mice per group. C-D) The protein levels of p-STAT3 and STAT3 (C) and the mRNA level of *Socs1* (D) in livers with the indicated AAV8 on day 3 after PHx. 3 mice per group. E) Western blots showing the levels of p-STAT3, STAT3, p-JAK2, JAK2 in CON and DKO livers at 12 hrs, day 1 and day 2 after PHx. F) The mRNA level of *Socs1* in CON and DKO livers on day 2 after PHx. 3 mice per group. G) IL-6 level in serum of CON and DKO mice at 12 hrs, day 1, day 2 after PHx. 4 mice per group for CON at 12 hrs and day 1, and 3 mice per group for others. H) Western blots showing the levels of p-STAT3, STAT3, p-JAK2, JAK2 in primary hepatocytes isolated from CON or DKO livers with or without 30 min treatment with IL-6. Data are presented as the mean ± SD. *: *P* <0.05. **: *P* <0.01.

To determine whether STAT3 mediates regulation of hepatocyte proliferation by executioner caspases, we overexpressed a constitutively active form of STAT3, STAT3C, in the DKO liver and evaluated the effect on regeneration after PHx. Overexpression of STAT3C in DKO livers dramatically upregulated the level of phosphorylated STAT3 and the expression of the target gene *Socs1* (Figure 7A & B). Overexpression of STAT3C in DKO livers rescued the reduced hepatocyte proliferation (Figure 7C & D) and impaired liver regeneration (Figure 7E), indicating that executioner caspases promote hepatocyte proliferation and liver regeneration by increasing STAT3 activity.

**Figure 7.**
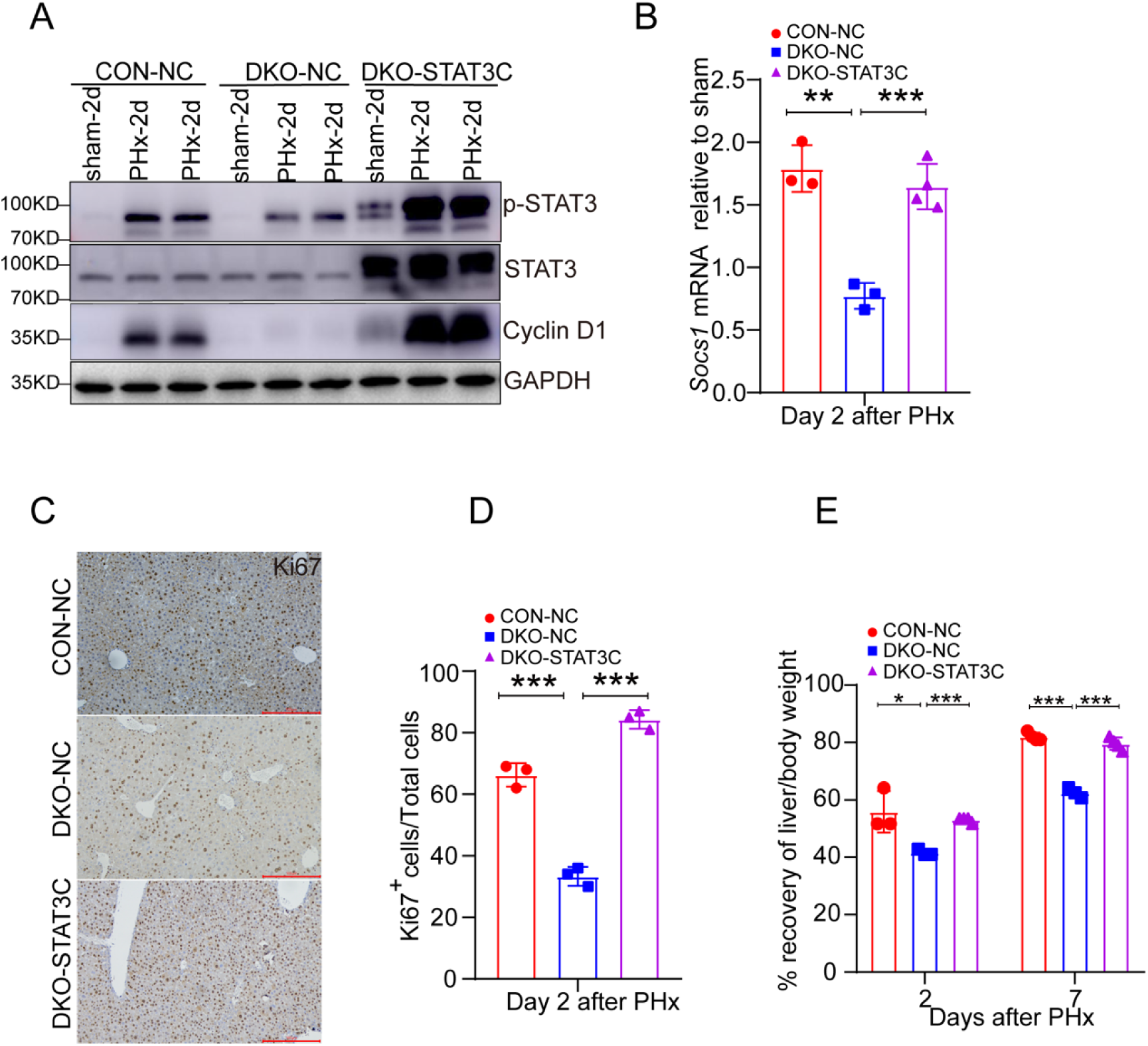
**Overexpression of the constitutively active form of STAT3 rescues the impaired hepatocyte proliferation and liver regeneration caused by loss of executioner caspases.** A) Western blots showing the levels of p-STAT3, STAT3 and Cyclin D1 in livers from CON mice infected with control AAV8 (CON-NC), DKO mice infected with control AAV8 (DKO-NC) and DKO mice infected with AAV8 carrying STAT3C (DKO-STAT3C) on day 2 after PHx. B) The mRNA level of *Socs1* in the indicated groups on day 2 after PHx. 3 mice per group. C-D) The representative images and quantification of Ki67 staining in the indicated groups on day 2 after PHx. Scale bar, 100 μm. 3 mice per group and 3 fields per mouse. E) The percentage of recovery of the liver/body weight ratio of the indicated mice on day 2 and day 7 after PHx. The average liver/body weight ratio of the sham group was considered as 100%. 3 mice for CON-NC and DKO-NC groups and 4 mice for DKO-STAT3C group. Data are presented as the mean ± SD. *: *P* <0.05. **: *P* <0.01. ***: *P* <0.001.

## Discussion

In this study, by generating transgenic mice carrying the mCasExpress lineage tracing system to label cells that have experienced ECA, we discover that ECA occurs at a low frequency in the homeostatic liver, but is dramatically upregulated during liver regeneration. ECA needs to be precisely controlled to promote proliferation of hepatocytes instead of killing them to achieve proper liver regeneration. Mechanistic studies reveal that ECA promotes hepatocyte proliferation through enhancing the activity of JAK/STAT3 signaling.

The mCasExpress sensor revealed enrichment of ECA in the pericentral region of the liver lobule under homeostasis. Previous work has shown that in homeostatic livers apoptotic bodies are mainly detected in the pericentral region^32, 33^. Wei et al. also reported that under homeostasis, hepatocytes in the pericentral region are gradually replaced by cells from the midlobular region^23^. These previous studies suggest that the turnover of hepatocytes involves apoptosis of the pericentral hepatocytes and generation of new hepatocytes from the midlobular region. Apoptosis can induce non-autonomous ECA and cell death^10, 34, 35^. In Drosophila pupal notum, the cells next to the dead cells experience ECA but survive through elevated ERK activity. Survival of these cells is necessary to ensure the epithelial integrity^10^. Thus, survival of hepatocytes with ECA in the pericentral region may be important for maintaining organ homeostasis through preventing excessive cell death in the process of removing old hepatocytes.

It has been reported that during regeneration after PHx, apoptosis rarely occurs in the first three days when hepatocytes are under rapid proliferation but is elevated in the termination phase^36, 37^, suggesting a role of apoptosis in terminating regenerative response. Interestingly, with the mCasExpress reporter we demonstrated that transient ECA took place more frequently in the first two days than in the termination phase. Lack of TUNEL^+^ staining in the livers in the first three days of regeneration suggests that the hepatocytes with transient ECA survived. Moreover, livers with caspase activity inhibited or with both *Casp3* and *Casp7* deleted exhibited reduced hepatocyte proliferation and impaired regeneration after PHx. The defective hepatocyte proliferation and regeneration in livers with *Casp3* or *Casp7* knocked out has also been reported by Li et al^8^. Their explanation for the phenomenon was that proliferation of hepatocytes during regeneration required executioner caspase-dependent secretion of prostaglandin E2 from the dying cells^8^. In addition, two studies recently reported that uptake of apoptotic extracellular vesicles by neutrophils or hepatocytes can promote liver regeneration^16, 17^. While these studies suggest the importance of dead cells in promoting liver regeneration, our data showed that the death of cells with ECA suppressed hepatocyte proliferation and liver regeneration after PHx, which indicates that the survival of cells with ECA is essential for regeneration. The enhanced proliferation in hepatocytes with ECA further supports the cell-autonomous effect of ECA on hepatocyte proliferation.

How does ECA promote hepatocyte proliferation? STAT3 is an important regulator of hepatocyte proliferation and liver regeneration. Mice with liver-specific knockout of *Stat3* displayed fewer proliferating hepatocytes and delayed regeneration^26,38^. We showed that inhibition of caspase activity or depletion of executioner caspases significantly suppressed STAT3 activation in the first two days of regeneration. Moreover, overexpression of the constitutively active form of STAT3, STAT3C, in livers lacking executioner caspases rescued the impaired hepatocyte proliferation and liver regeneration after PHx. These data support that executioner caspases promote regenerative proliferation through enhancing STAT3 activity. Upon liver injury, STAT3 can be activated by IL-6 secreted from macrophages and hepatocytes^20, 30, 39^. IL-6 binding to its receptor results in recruitment and phosphorylation of JAK, which in return lead to STAT3 phosphorylation and activation^31^. We found that primary hepatocytes lacking executioner caspases displayed reduced phosphorylation of JAK2 and STAT3 upon IL-6 treatment in vitro, suggesting that executioner caspases regulate STAT3 activity downstream of IL-6 and upstream of JAK. Several proteins in JAK/STAT3 pathway have been reported as substrates of caspases, yet the caspase-mediated cleavage of these proteins leads to inhibition of the pathway activity. For example, STAT3 is cleaved in a caspase-dependent manner in cancer cells treated with staurosporine, resulting in inactivation of STAT3^40^. In liver, activated caspase-3 induced by CD95L can cleave gp130, a subunit of IL-6 receptor complex, thereby suppressing STAT3 activation^41, 42^. Activation of JAK/STAT3 signaling by executioner caspases has rarely been reported. In cancer cells, low-level caspase-3 activation can cause DNA damage by activating endonuclease G and caspase-dependent DNase. The DNA damage then leads to phosphorylation of STAT3 via activation of Src^29, 43^. However, the mechanism underlying caspase-induced STAT3 activation during liver regeneration may be different from that in cancer cells, as little DNA damage was detected in regenerating livers. Moreover, our data suggest that executioner caspases regulate STAT3 upstream of JAK but Src has been reported to activate STAT3 in parallel to JAK^44^.

In summary, our work identifies executioner caspases as positive regulators of JAK/STAT3 signaling to promote hepatocyte proliferation during liver regeneration. The ECA in hepatocytes needs to be tightly controlled. Both too little and too much ECA impedes regeneration.

## Materials and methods

### Mice

All mouse experiments were performed in accordance with the protocol approved by the Institutional Animal Care and Use Committee at School of Basic Medical Sciences, Shandong University (ECSBMSSDU2023-2-141). Generation of the transgenic mice carrying *CAG-FSF-ZsGreen* and *CAG-LSL-rtTA-TRE-Lyn11-NES-DEVD-FLP* were done by GemPharmatech Co., Ltd (Nanjing, China) using CRISPR-Cas9 technology. *Sox2-Cre* (Stain# 008454), *tetO-Cre* (Strain# 006234) and *CAG-LSL-tdTomato* mice (Strain# 007909) were obtained from the Jackson Laboratory (Bar Harbor, Maine, US). *CAG-Cre* (Strain# T050269), *Alb-Cre* (Strain# T003814), *Casp3^flox/flox^* (Strain# T005781) and *Casp7^flox/flox^* (Strain# T005784) were obtained from GemPharmatech. *CAG-rtTA* mice (Strain# C001185) were obtained from Cyagen Biosciences (Suzhou, China). All the mice were housed in a specific-pathogen-free facility at 22-26 °C and 40-70% humidity, with a 12/12 h light-dark cycle. To induce expression of *Lyn11-NES-DEVD-FLP*, doxycycline (DOX; Sangon Biotech Cat# A600889-0025) was dissolved in 0.9% NaCl and administered to mice through tail veins at 5 mg per kg mouse body weight.

### Genotyping

For genotyping, mouse tails (∼ 0.1 cm) were incubated with Proteinase K (Spark jade, Cat# AA1906) overnight at 55 °C, followed by 10 min incubation at 100 °C to inactivate Proteinase K. After centrifugation at 12000 rpm for 10 min, the supernatant containing genomic DNA was applied to PCR. The primers were synthesized by Tsingke Biotech (Beijing, China). The primers used for genotyping are listed in Supplementary table S1.

### Liver injury

70% PHx was performed as reported in literature^45^. In brief, 8-week-old male mice were anaesthetized with isoflurane and oxygen flow. The abdominal skin and muscle were incised to expose the liver. The left lateral and median hepatic lobes were ligated and removed. After closing the abdominal cavity, betadine was applied to the suture and the mice were kept in a 37 °C incubator to recover. For CCl_4_-induced liver injury model, 10% CCl_4_ (dissolved in corn oil) were administered intraperitoneally into 8-week-old male mice at a dose of 10 μl per gram body weight. The mice were euthanized at the expected time points and the liver weight and body weight were measured. The serum AST and ALT were determined by Kingmed Diagnostics (Guangzhou, China).

### AAV injection

AAV8-CAG-XIAP1, AAV8-CAG-p35, AAV8-CAG-FSF-tBid, AAV8-CAG-STAT3C and the control AAV8 virus were provided by GeneChem (Shanghai, China). The virus was injected into 6-week-old mice through tail veins at a dose of 3×10^11^ vg per mouse.

### Histology and immunohistochemistry

The liver tissues were fixed with 4% paraformaldehyde (ServiceBio, Cat# G1101) for 48 hours, and embedded in paraffin (ServiceBio). The embedded tissues were sectioned and stained with hematoxylin and eosin. For immunohistochemistry, the tissue sections were subjected to antigen retrieval with citric acid antigen retrieval buffer (PH6.0) followed by three times wash with PBS. Activity of endogenous peroxidase was blocked by incubation with 3% hydrogen peroxide for 30 min. The sections were then incubated with 3% BSA for 30 minutes at room temperature. Primary antibodies were incubated overnight at 4 °C. Secondary antibodies were incubated at 37 °C for 50 minutes. The signals were detected using DAB (ORIGENE, Cat# ZLI-9108). The images were taken on an upright microscope (Olympus). The antibodies used are listed in Supplementary table S2.

### Immunofluorescent staining

Liver tissues were fixed in 4% paraformaldehyde, equilibrated to 30% sucrose and frozen in O.C.T. compound. 6 μm sections were cut, washed with PBS and blocked with 5% goat serum. The sections were then incubated with primary antibody at 4 °C overnight and secondary antibody at 37 °C for 1 hr. The images were captured on an upright fluorescent microscope (Olympus). All the antibodies used for staining are listed in Supplementary table S2. To detect apoptotic cells, TUNEL assays were performed using TUNEL BrightRed Apoptosis Detection Kit (Vazyme, Cat# A113-03) according to the manufacturer’s protocol. For picrosirius red staining, the liver sections were stained according to the published protocol^46^ and imaged under polarized light using a NIKON Eclipse ci microscope.

### Quantification of zonal location

Quantification of zonal location was done according to the literature^23^. Glutamine synthase (GS) was stained to mark the central veins. The position index (P.I.) of each ZsGreen^+^ cell was calculated based on its distance to the closest CV (x), the distance to the closest PV (y) and the distance between the CV and PV (z) using the formula P.I. = (x^2^ + z^2^ - y^2^)/(2z^2^) (Figure 1G). Cells with P.I. less than 0.33 were considered in zone 3. Cells with P.I. between 0.33 and 0.66 were considered in zone 2. Cells with P.I. more than 0.66 were considered in zone 1.

### Isolation of primary hepatocytes and treatment with IL-6

Hepatocytes were isolated by two-step collagenase perfusion modified from a published protocol^47^. The liver was perfused through the portal vein with the perfusion buffer (HBSS containing 0.5 mM EGTA, PH7.4) followed by the digestion buffer (HBSS containing 5 mM CaCl_2_, 0.1 mg/ml Collagenase IV and 10 mM HEPES, PH 7.4). The cell suspension was centrifuged at 50 g for 3 min. The pellets were resuspended in DMEM/F12 (Gibco, Cat# 8123375) supplemented with 10% FBS and 100 U/ml penicillin and 100 μg/ml streptomycin and seeded on collagen-coated 6-well plates at a density of 3.5×10^5^ cells/well. After 16-hour culture, the hepatocytes were serum-starved for 16 hours and then treated with human recombinant IL-6 (Peprotech, Cat# 200-06) for 30 min.

### RNA extraction and quantitative RT-PCR

Total RNA was extracted using FastPure cell/tissue total RNA isolation kit V2 (Vazyme, Cat# R112-01) and converted to cDNA using HiScript III RT SuperMix for qPCR kits (Vazyme, Cat# R323-01). Quantitative PCR was performed using ChamQ SYBR Color qPCR Master Mix (Vazyme, Cat# Q411-02). The primers used are listed in the Supplementary table S1.

### Western blot

The frozen liver tissues were homogenized and lysed in RIPA buffer (Sigma, Cat# SLBZ0792). Protein concentrations were measured with the BCA assay kit (Vazyme, Cat# E112-02). 20 µg proteins were applied to SDS-PAGE then transferred to PVDF membranes. The membranes were incubated with primary antibody (1:1000) overnight at 4°C and then with the secondary antibody (1:2000) at room temperature for 1h. Proteins were detected using Chemiluminescent Substrates (Spark jade, Cat# ED0015-C) and Tanon 5200 Multi Chemiluminescence imager (Tanon Science & Technology Co., Shanghai, China). The antibodies used are listed in the Supplementary table S2.

### Statistical analysis

Data are presented as the mean ± standard deviation (SD). Statistical significance was determined using t-test for two-sample comparison or one-way ANOVA for comparing three or more samples. The Tukey test was used to derive adjusted *P*-value for multiple comparisons. *P*<0.05 was considered as statistically significant. The assumption of equal variance was validated by F-test. Statistical analyses were performed using GraphPad Prism version 8 (GraphPad Software). The sample sizes were chosen empirically based on the observed effects and previous reports. The sample size for each experiment is listed in the figure legends. When collecting and analyzing data of immunohistochemistry and immunofluorescent staining, the investigators were blinded to the group allocation.

### Data availability statement

The original Western blots are provided. All the other raw data supporting the findings of this study are available from the corresponding author upon request.

## Acknowledgement

We thank Translational Medicine Core Facility of Shandong University and the School of Basic Medical Sciences Core Facility of Shandong University for technical support.

## Author contribution

ZC and GS designed the experiments. ZC, LQ, KL, XH and EL performed the experiments. MW, BJ, YZ, HH and QL provided resources and technical supports. ZC, CS, YG and GS prepared the manuscript.

## Ethics approval statement

All mouse experiments were performed in accordance with the protocol approved by the Institutional Animal Care and Use Committee at School of Basic Medical Sciences, Shandong University (ECSBMSSDU2023-2-141).

## Reference

1. Sun, G. Death and survival from executioner caspase activation. Semin Cell Dev Biol (2023).

2. Yuan, J. & Ofengeim, D. A guide to cell death pathways. Nat Rev Mol Cell Biol (2023).

3. Chera, S. et al. Apoptotic cells provide an unexpected source of Wnt3 signaling to drive hydra head regeneration. Dev Cell 17, 279–289 (2009).

4. Wells, B.S., Yoshida, E. & Johnston, L.A. Compensatory proliferation in Drosophila imaginal discs requires Dronc-dependent p53 activity. Curr Biol 16, 1606–1615 (2006).

5. Gauron, C. et al. Sustained production of ROS triggers compensatory proliferation and is required for regeneration to proceed. Sci Rep 3, 2084 (2013).

6. Tseng, A.S., Adams, D.S., Qiu, D., Koustubhan, P. & Levin, M. Apoptosis is required during early stages of tail regeneration in Xenopus laevis. Dev Biol 301, 62–69 (2007).

7. Wang, H. et al. Turning terminally differentiated skeletal muscle cells into regenerative progenitors. Nat Commun 6, 7916 (2015).

8. Li, F. et al. Apoptotic cells activate the “phoenix rising” pathway to promote wound healing and tissue regeneration. Sci Signal 3, ra13 (2010).

9. Ankawa, R. et al. Apoptotic cells represent a dynamic stem cell niche governing proliferation and tissue regeneration. Dev Cell 56, 1900–1916 e1905 (2021).

10. Valon, L. et al. Robustness of epithelial sealing is an emerging property of local ERK feedback driven by cell elimination. Dev Cell 56, 1700–1711 e1708 (2021).

11. Medina, C.B. et al. Metabolites released from apoptotic cells act as tissue messengers. Nature 580, 130–135 (2020).

12. Sebbagh, M. et al. Caspase-3-mediated cleavage of ROCK I induces MLC phosphorylation and apoptotic membrane blebbing. Nat Cell Biol 3, 346–352 (2001).

13. Coleman, M.L. et al. Membrane blebbing during apoptosis results from caspase-mediated activation of ROCK I. Nat Cell Biol 3, 339–345 (2001).

14. Tixeira, R. et al. ROCK1 but not LIMK1 or PAK2 is a key regulator of apoptotic membrane blebbing and cell disassembly. Cell Death Differ 27, 102–116 (2020).

15. Brock, C.K. et al. Stem cell proliferation is induced by apoptotic bodies from dying cells during epithelial tissue maintenance. Nat Commun 10, 1044 (2019).

16. Brandel, V. et al. Hepatectomy-induced apoptotic extracellular vesicles stimulate neutrophils to secrete regenerative growth factors. J Hepatol 77, 1619–1630 (2022).

17. Sui, B. et al. Apoptotic Vesicular Metabolism Contributes to Organelle Assembly and Safeguards Liver Homeostasis and Regeneration. Gastroenterology 167, 343–356 (2024).

18. Tang, H.L. et al. Cell survival, DNA damage, and oncogenic transformation after a transient and reversible apoptotic response. Mol Biol Cell 23, 2240–2252 (2012).

19. Sun, G., Ding, X.A., Argaw, Y., Guo, X. & Montell, D.J. Akt1 and dCIZ1 promote cell survival from apoptotic caspase activation during regeneration and oncogenic overgrowth. Nat Commun 11, 5726 (2020).

20. Michalopoulos, G.K. & Bhushan, B. Liver regeneration: biological and pathological mechanisms and implications. Nat Rev Gastroenterol Hepatol 18, 40–55 (2021).

21. Ekert, P.G., Silke, J. & Vaux, D.L. Caspase inhibitors. Cell Death Differ 6, 1081–1086 (1999).

22. Ben-Moshe, S. & Itzkovitz, S. Spatial heterogeneity in the mammalian liver. Nat Rev Gastroenterol Hepatol 16, 395–410 (2019).

23. Wei, Y. et al. Liver homeostasis is maintained by midlobular zone 2 hepatocytes. Science 371 (2021).

24. Wei, M.C. et al. tBID, a membrane-targeted death ligand, oligomerizes BAK to release cytochrome c. Genes Dev 14, 2060–2071 (2000).

25. Flores-Romero, H. et al. BCL-2-family protein tBID can act as a BAX-like effector of apoptosis. EMBO J 41, e108690 (2022).

26. Li, W., Liang, X., Kellendonk, C., Poli, V. & Taub, R. STAT3 contributes to the mitogenic response of hepatocytes during liver regeneration. J Biol Chem 277, 28411–28417 (2002).

27. Haga, S. et al. Compensatory recovery of liver mass by Akt-mediated hepatocellular hypertrophy in liver-specific STAT3-deficient mice. J Hepatol 43, 799–807 (2005).

28. Huda, K.A. et al. Ex vivo adenoviral gene transfer of constitutively activated STAT3 reduces post-transplant liver injury and promotes regeneration in a 20% rat partial liver transplant model. Transpl Int 19, 415–423 (2006).

29. Zhu, C., Fan, F., Li, C.Y., Xiong, Y. & Liu, X. Caspase-3 promotes oncogene-induced malignant transformation via EndoG-dependent Src-STAT3 phosphorylation. Cell Death Dis 15, 486 (2024).

30. Cressman, D.E. et al. Liver failure and defective hepatocyte regeneration in interleukin-6-deficient mice. Science 274, 1379–1383 (1996).

31. Philips, R.L. et al. The JAK-STAT pathway at 30: Much learned, much more to do. Cell 185, 3857–3876 (2022).

32. Benedetti, A., Jezequel, A.M. & Orlandi, F. Preferential distribution of apoptotic bodies in acinar zone 3 of normal human and rat liver. J Hepatol 7, 319–324 (1988).

33. Benedetti, A., Jezequel, A.M. & Orlandi, F. A quantitative evaluation of apoptotic bodies in rat liver. Liver 8, 172–177 (1988).

34. Perez-Garijo, A., Fuchs, Y. & Steller, H. Apoptotic cells can induce non-autonomous apoptosis through the TNF pathway. Elife 2, e01004 (2013).

35. Herrera, S.C., Martin, R. & Morata, G. Tissue homeostasis in the wing disc of Drosophila melanogaster: immediate response to massive damage during development. PLoS Genet 9, e1003446 (2013).

36. Sakamoto, T. et al. Mitosis and apoptosis in the liver of interleukin-6-deficient mice after partial hepatectomy. Hepatology 29, 403–411 (1999).

37. Ozaki, M. Cellular and molecular mechanisms of liver regeneration: Proliferation, growth, death and protection of hepatocytes. Semin Cell Dev Biol 100, 62–73 (2020).

38. Haga, S. et al. Stat3 protects against Fas-induced liver injury by redox-dependent and -independent mechanisms. J Clin Invest 112, 989–998 (2003).

39. Norris, C.A. et al. Synthesis of IL-6 by hepatocytes is a normal response to common hepatic stimuli. PLoS One 9, e96053 (2014).

40. Darnowski, J.W. et al. Stat3 cleavage by caspases: impact on full-length Stat3 expression, fragment formation, and transcriptional activity. J Biol Chem 281, 17707–17717 (2006).

41. Graf, D. et al. Bile acids inhibit interleukin-6 signaling via gp130 receptor-dependent and -independent pathways in rat liver. Hepatology 44, 1206–1217 (2006).

42. Graf, D., Haselow, K., Munks, I., Bode, J.G. & Haussinger, D. Caspase-mediated cleavage of the signal-transducing IL-6 receptor subunit gp130. Arch Biochem Biophys 477, 330–338 (2008).

43. Liu, X. et al. Self-inflicted DNA double-strand breaks sustain tumorigenicity and stemness of cancer cells. Cell Res 27, 764–783 (2017).

44. Garcia, R. et al. Constitutive activation of Stat3 by the Src and JAK tyrosine kinases participates in growth regulation of human breast carcinoma cells. Oncogene 20, 2499–2513 (2001).

45. Pu, W. et al. Mfsd2a+ hepatocytes repopulate the liver during injury and regeneration. Nat Commun 7, 13369 (2016).

46. Rittie, L. Method for Picrosirius Red-Polarization Detection of Collagen Fibers in Tissue Sections. Methods Mol Biol 1627, 395–407 (2017).

47. Charni-Natan, M. & Goldstein, I. Protocol for Primary Mouse Hepatocyte Isolation. STAR Protoc 1, 100086 (2020).

