## Supplementary figures and tables for "Survival of hepatocytes from executioner caspase activation promotes liver regeneration by enhancing JAK/STAT3 activity"

Cao et al.

This file contains the following contents:

Supplementary figure S1-S11 and their legends

Supplementary table S1-S2

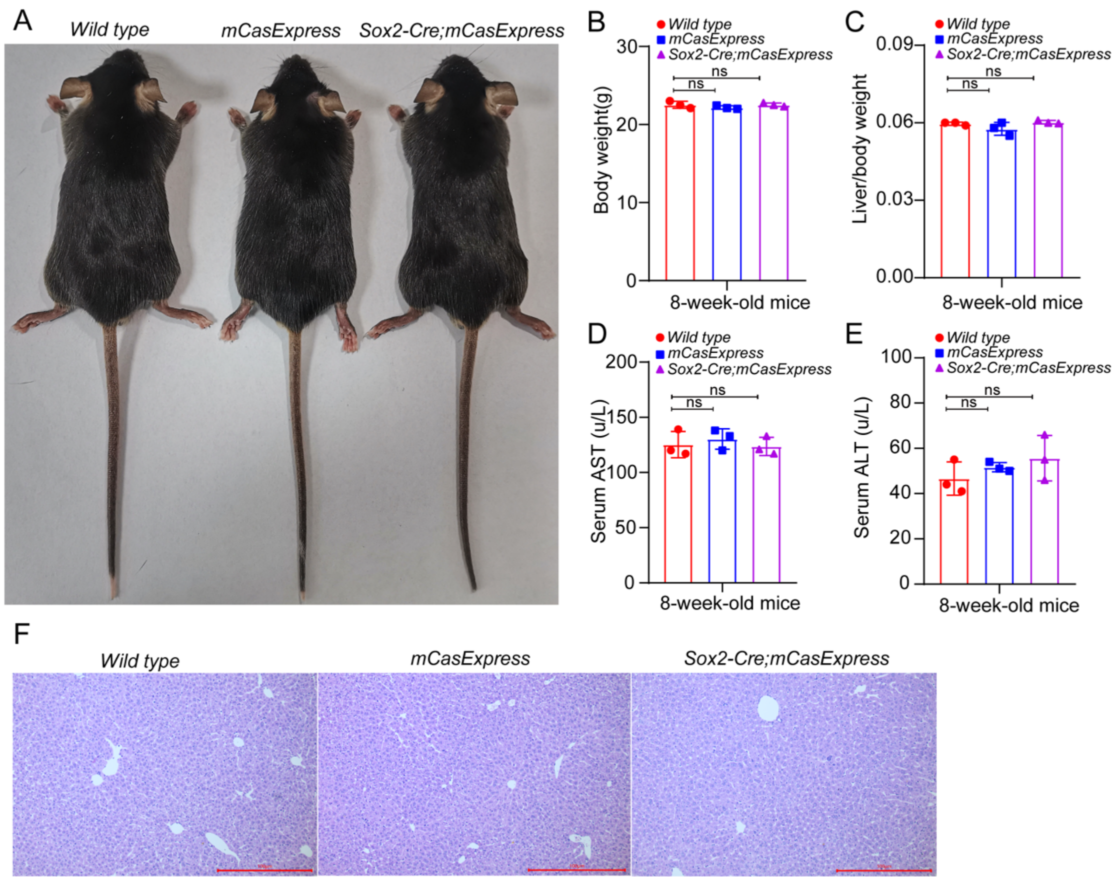


**Supplementary figure S1. The *mCasExpress* and *Sox2-Cre; mCasExpress* mice have body and liver morphology comparable to the wild type mice.**

A) Pictures of the 8-week-old mice with the indicated genotypes. B-E) The body weight (B), the ratio between liver weight and body weight (C), the serum AST (D) and the serum ALT (E) of the *wild type*, *mCasExpress*, *Sox2-Cre; mCasExpress* mice. 3 mice per group. Data are presented as the mean ± SD. F) The representative images of H & E staining of livers from the indicated mice. Scale bar: 100 µm. ns: no significance.

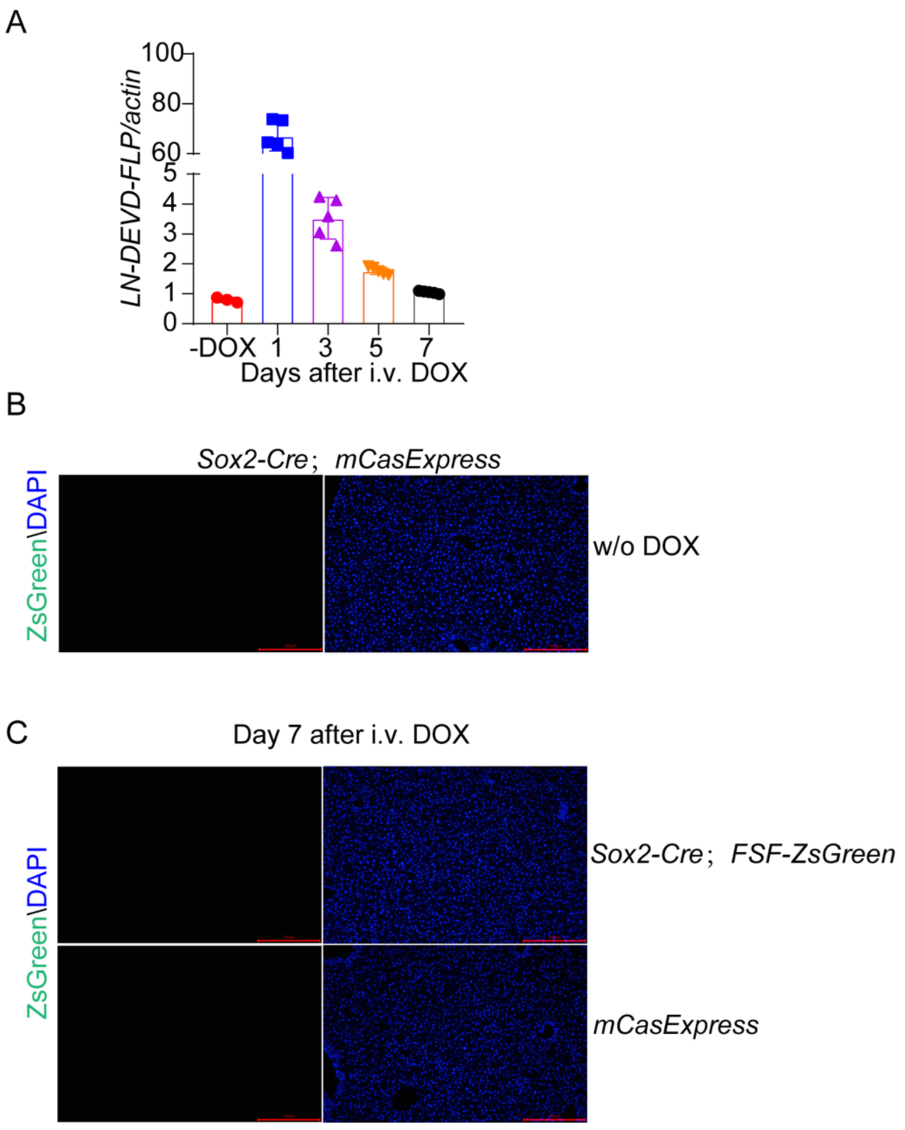


**Supplementary figure S2. Expression of ZsGreen requires the presence of DOX, Cre and FLP.**

A) The relative mRNA level of *LN-DEVD-FLP* before (-DOX) and after DOX injection. 3 mice in -DOX group and 5 mice in all the other groups. Data are presented as the mean ± SD. B) The representative images of the livers from *Sox2-Cre; mCasExpress* mice without DOX injection. Scale bar: 100 μm. C) The representative images of the livers from *mCasExpress* mice and *Sox2-Cre; FSF-ZsGreen* mice on day 7 after DOX injection. Scale bar: 100 μm. i.v.: intravenous injection.

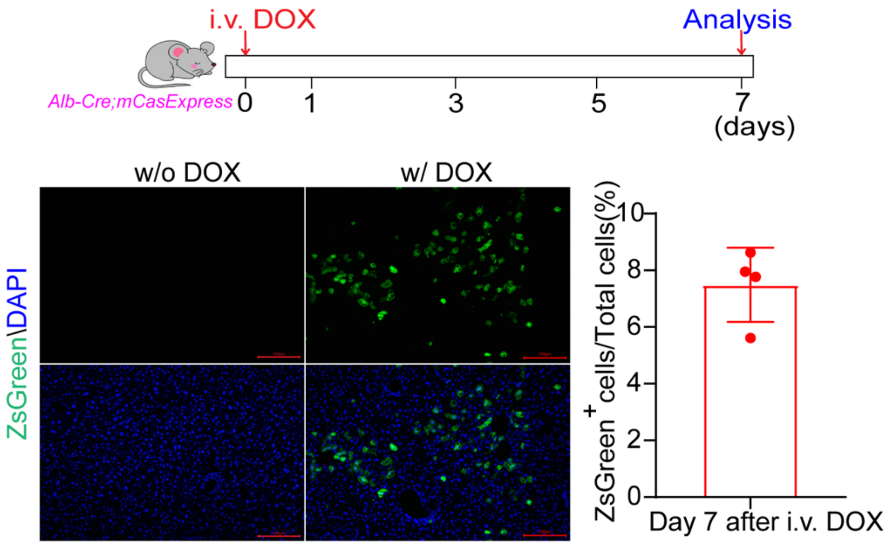


**Supplementary figure S3. ECA occurs at low frequency in homeostatic livers.**

The representative images and quantification of the ZsGreen$^{+}$ cells in the *Alb-Cre; mCasExpress* livers without DOX injection (w/o DOX) or on day 7 after DOX injection (w/ DOX). Scale bar: 100 µm. 4 mice. Data are presented as the mean ± SD.

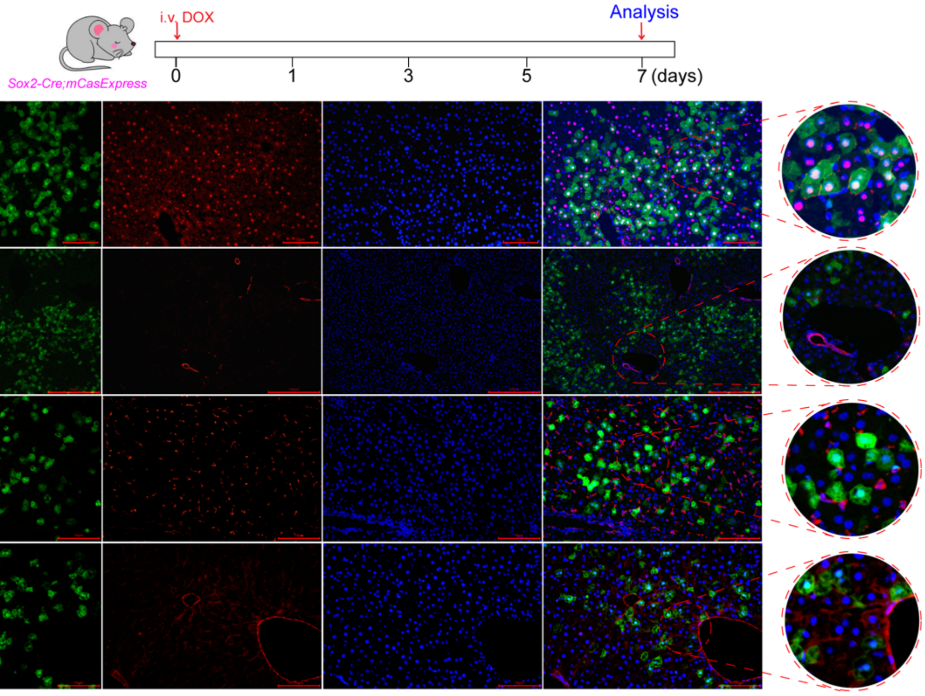


**Supplementary figure S4. ECA occurs only in hepatocytes.**

The representative images of livers with staining of the hepatocyte marker HNF4α, cholangiocyte marker CK19, the Kupffer cell marker CD68 or the endothelial cell marker CD31. Scale bar: 50 μm for HNF4α, CD68 and CD31 images and 100 μm for CK19 images.

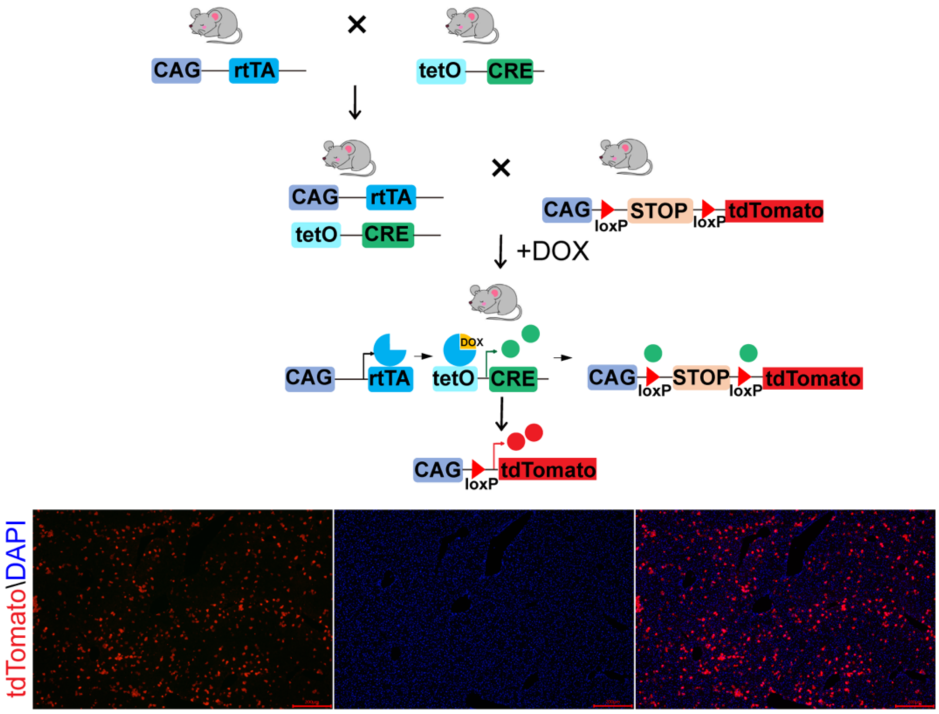


**Supplementary figure S5. Response to DOX is similar across all zones in the liver lobule.**

The upper part shows the mating strategy and the lower are the representative images of tdTomato expression in the livers from the *CAG-rtTA; tetO-Cre; LSL-tdTomato* mice on day 7 after DOX injection. Scale bar: 200 μm.

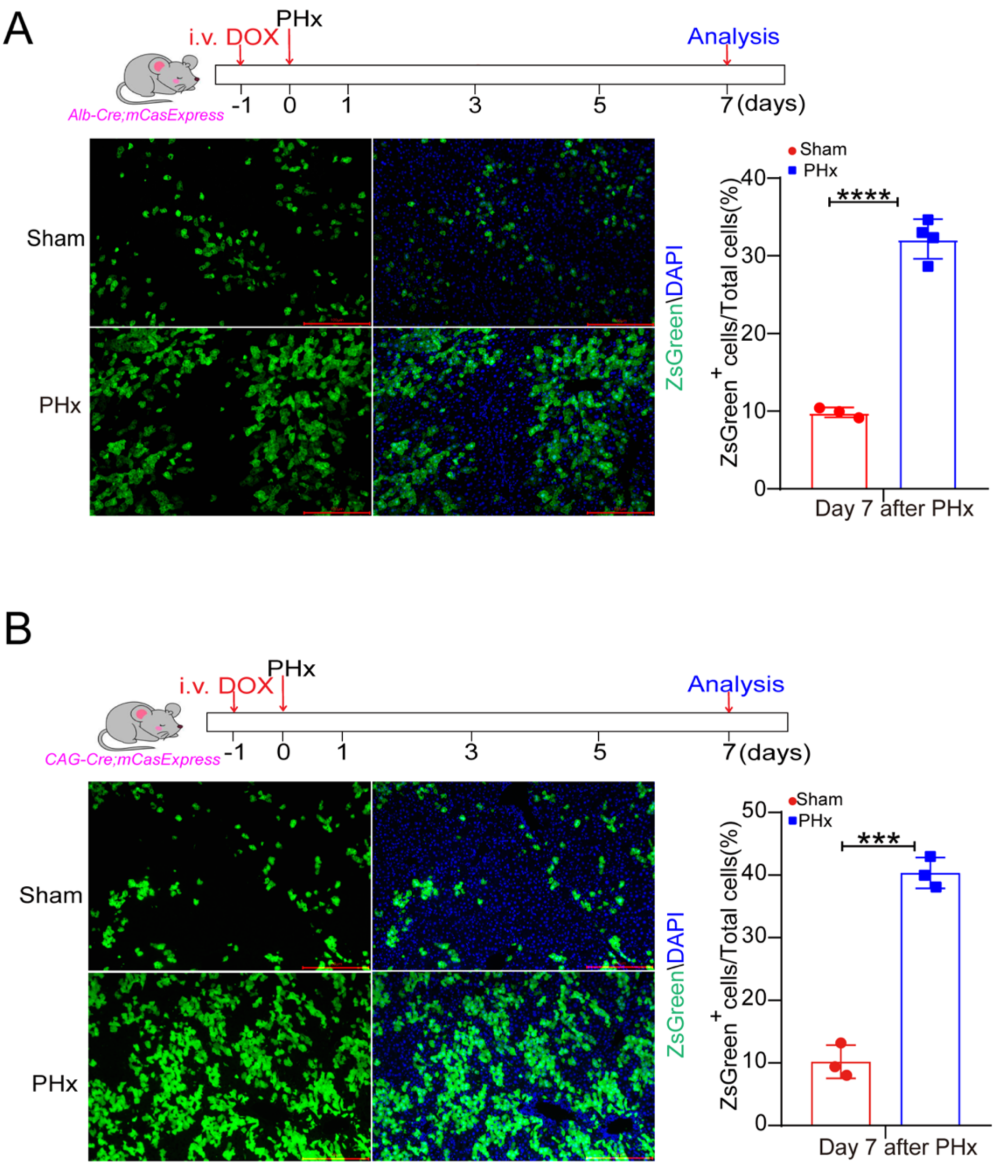


**Supplementary figure S6. ECA is dramatically elevated during regeneration after PHx.**

The representative images and quantification of ZsGreen$^{+}$ cells in *Alb-Cre; mCasExpress* livers (A) and *CAG-Cre; mCasExpress* livers (B) on day 7 after PHx. Scale bar: 100 µm. 4 mice in the PHx group of *Alb-Cre; mCasExpress* and 3 mice per group for all the others. 3 fields per mouse. Data are presented as the mean ± SD. ***: *P* <0.001. ****: *P* <0.0001.

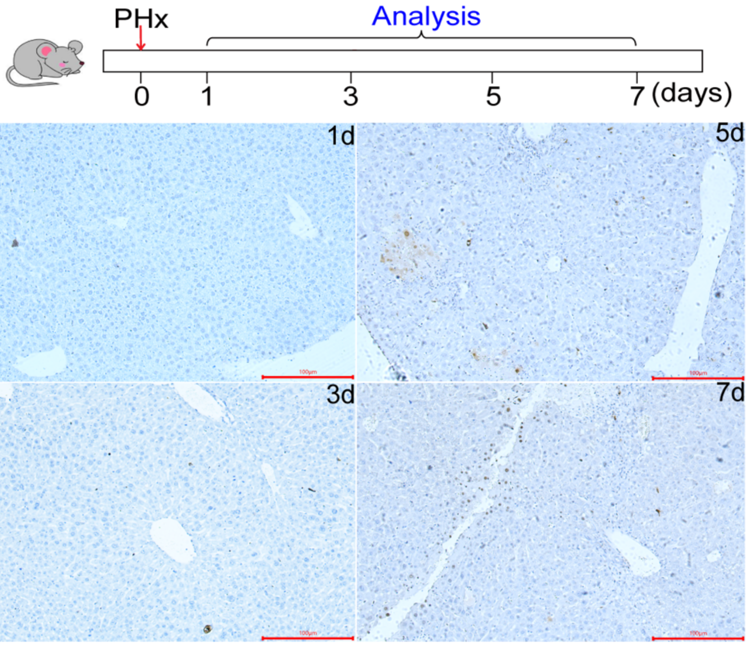


**Supplementary figure S7. Apoptotic cell death during regeneration after PHx.**

The representative images of TUNEL staining on livers collected on day 1, 3, 5, 7 after PHx. Scale bar: 100 µm.

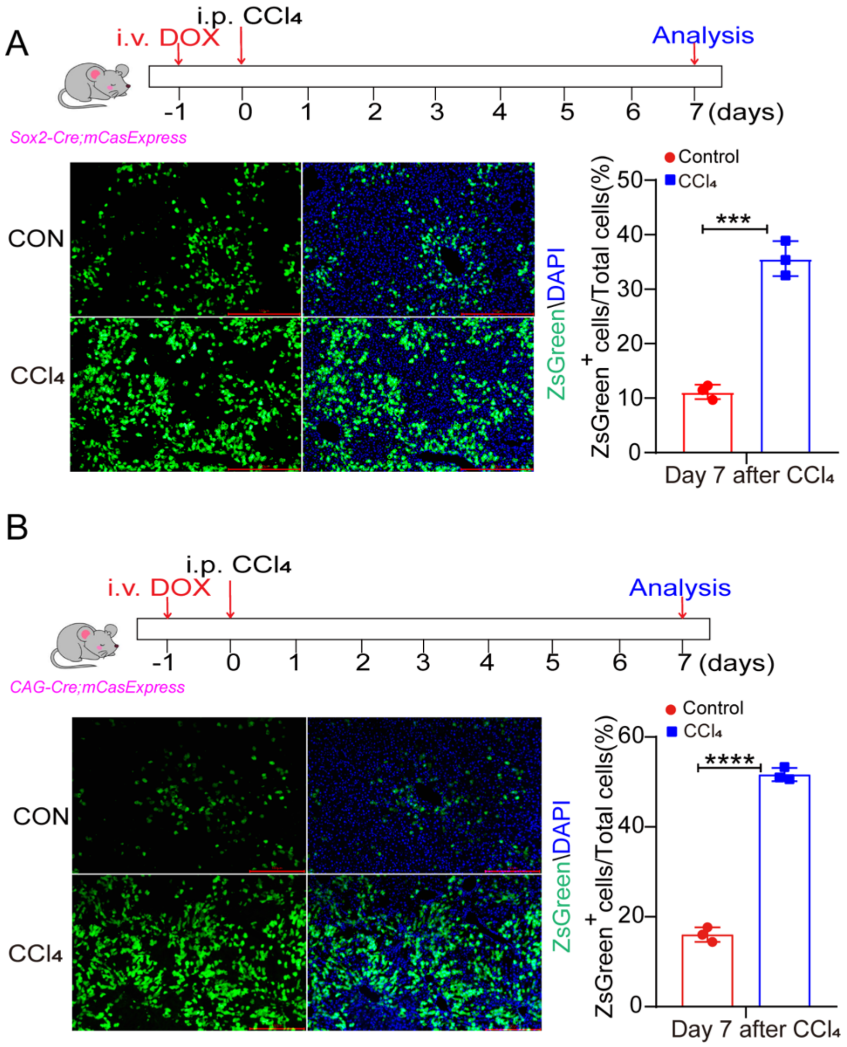


**Supplementary figure S8. ECA is dramatically increased in livers during regeneration after acute $CCl_4$ injury.**

The representative images and quantification of the ZsGreen$^+$ cells in livers from *Sox2-Cre; mCasExpress* mice (A) or *CAG-Cre; mCasExpress* mice (B) on day 7 after injection of $CCl_4$ or corn oil (CON). Scale bar: 100 µm. 3 mice per group and 3 fields per mouse. i.v.: intravenous injection. i.p. intraperitoneal injection. Data are presented as the mean ± SD. ***: *P* <0.001. ****: *P* <0.0001.

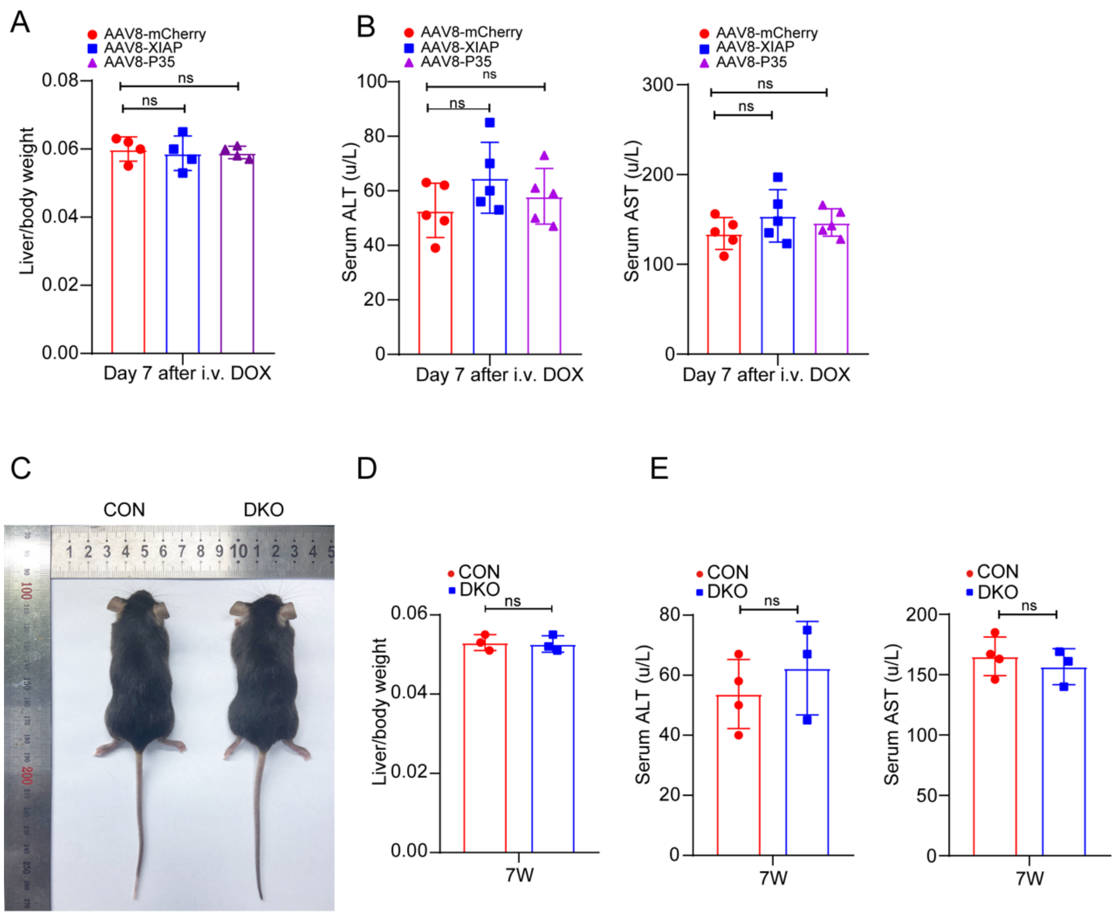


**Supplementary figure S9. Inhibition of executioner caspase activity or expression does not affect liver weight or function.**

A-B) The ratio between liver weight and body weight (4 mice per group), serum AST and ALT (5 mice per group) in mice injected with the indicated AAV8. C) The image of CON and DKO mice. D-E) The liver/body weight ratio (D), serum ALT and AST levels (E) of the 7-week-old male CON and DKO. 3 or 4 mice per group. Data are presented as the mean ± SD. ns: no significance.

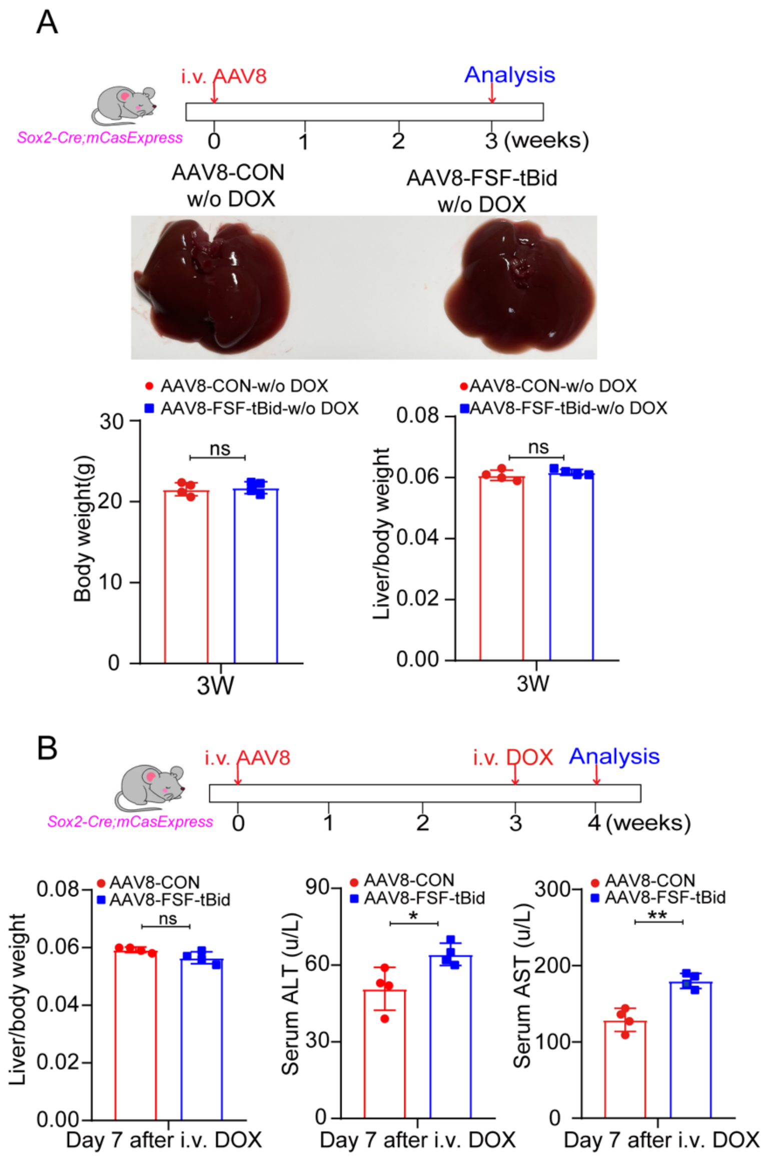


**Supplementary figure S10. Ablating cells with ECA in homeostatic livers mildly affects liver function.**

A) Without DOX injection, AAV8-FSF-tBid showed little effect on liver morphology, body weight and liver/body weight ratio. 4 mice per group. B) Mice administered with AAV8-FSF-tBid exhibited similar liver/body weight ratio but mildly increased serum AST and ALT. 4 mice per group. i.v.: intravenous injection. Data are presented as the mean ± SD. *: $P$ <0.05. **: $P$ <0.01. ns: no significance.

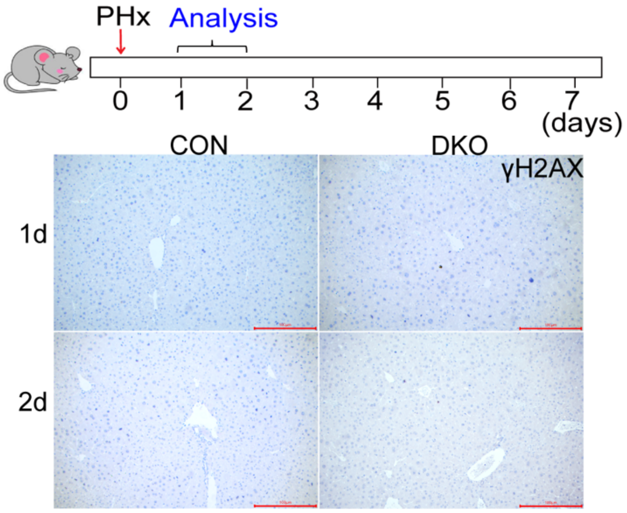


**Supplementary figure S11. PHx does not induce DNA damage in livers.**

γH2AX staining in CON or DKO mice on day 1 and 2 after PHx. Scale bar: 100 μm.

**Supplementary table S1. The list of primers.**

| **qPCR** | | |
|---|---|---|
| **Gene name** | **Forward primer** | **Reverse primer** |
| *Ccnd1* | AGGCGGATGAGAACAAGCAG | AGAAAGTGCGTTGTGCGGTA |
| *Ccne1* | CCTTTCAGTCCGCTCCAGAA | GGATGAAAGAGCAGGGGTCC |
| *Socs1* | CTGCGGCTTCTATTGGGGAC | AAAAGGCAGTCGAAGGTCTCG |
| *FLP* | TGGTGTACCTGGACGAGTTCCTG | CTGCTTGTTGCTGCTGCTGTTG |
| *Actin* | GTGCTATGTTGCTCTAGACTTCG | ATGCCACAGGATTCCATACC |
| **Genotyping** | | |
| **Gene name** | **Forward primer** | **Reverse primer** |
| *LN-DEVD-FLP* | ATCATCCCTTACAACGGCCA | CCGGTCCTGTTCACTCTCTT |
| *FSF-ZsGreen* | AGATGACCATGAAGTACCGCA | CTCCCAGTTGTCGGTCATCT |
| *Sox2-Cre* | CCCGCAGAACCTGAAGATG | GACCCGGCAAAACAGGTAG |
| *CAG-Cre* | TTCGGCTTCTGGCGTGTGA | CTGACTTCATCAGAGGTGGCATC |
| *Alb-Cre* | TGGATGCCACCTCTGATGAAGTC | TCCTGGCATCTGTCAGAGTTCTCC |
| *Alb-Cre WT* | CAGCAAAACCTGGCTGTGGATC | ATGAGCCACCATGTGGGTGTC |
| *tetO-Cre* | GCGGTCTGGCAGTAAAAA CTA TC | GTGAAACAGCATTGCTGTCAC TT |

| | | |
|---|---|---|
| *tdTomato* | CGGCATGGACGAGCTGTACAAG | TCAGCAAACACAGTGCACACCAC |
| *tdTomato WT* | CCCAAAGTCGCTCTGAGTTGTTA | TCGGGTGAGCATGTCTTTAATCT |
| *Casp3^flox^* | GCATCGCATTGTCTGAGTAGGTG | TACTTGGTCCCGAGTAAGTGGAAC |
| *Casp3 WT* | CAGCAAAACCTGGCTGTGGATC | ATGAGCCACCATGTGGGTGTC |
| *Casp7^flox^* | CATGAAAGGTCTGGGATTGTG | CCGCCCCTGTATGTTTTG |
| *rtTA* | AAAGTCGCTCTGAGTTGTTAT | GCGAAGAGTTTGTCCTCAACC |
| *rtTA WT* | AAAGTCGCTCTGAGTTGTTAT | GGAGCGGGAGAAATGGATATG |
| *H11* | CAGCAAAACCTGGCTGTGGATC | ATGAGCCACCATGTGGGTGTC |
| *ROSA26* | CCCAAAGTCGCTCTGAGTTGTTA | TCGGGTGAGCATGTCTTTAATCT |

**Supplementary table S2. The list of antibodies.**

| **Antibody** | **Company** | **Catalog No.** |
|---|---|---|
| Anti-CK19 | Servicebio | GB12197 |
| Anti-CD31 | Servicebio | GB11315 |
| Anti-CD68 | Cell Signaling Technology | 97778 |
| Anti-Glutamine Synthetase | Abcam | Ab176562 |
| Anti-Ki67(IF) | Abcam | Ab15580 |
| Anti-Ki67(IHC) | Cell Signaling Technology | 12202 |
| Anti-GAPDH | Proteintech | 60004-1-1g |
| Anti-STAT3 | Cell Signaling Technology | 9139 |
| Anti-phosphor-STAT3 | Cell Signaling Technology | 9145 |
| Anti-Cyclin D1 | Abways | CY5404 |
| Anti-JAK2 | Santa Cruz Biotechnology | sc-390539 |
| Anti-phosphor-JAK2 | Cell Signaling Technology | 3771 |
| HRP-goat anti-mouse | ORIGENE | ZB-2305 |
| HRP-goat anti-rabbit | ORIGENE | ZB-2301 |
